# Motility induced fracture reveals a ductile to brittle crossover in the epithelial tissues of a simple animal

**DOI:** 10.1101/676866

**Authors:** Vivek N. Prakash, Matthew S. Bull, Manu Prakash

## Abstract

Animals are characterized by their movement, and their tissues are continuously subjected to dynamic force loading while they crawl, walk, run or swim^1^. Tissue mechanics fundamentally determine the ecological niches that can be endured by a living organism^2^. While epithelial tissues provide an important barrier function in animals, they are subjected to extreme strains during day to day physiological activities, such as breathing^1^, feeding^3^, and defense response^4^. However, failure or inability to withstand to these extreme strains can result in epithelial fractures^5, 6^ and associated diseases^7, 8^. From a materials science perspective, how properties of living cells and their interactions prescribe larger scale tissue rheology and adaptive response in dynamic force landscapes remains an important frontier^9^. Motivated by pushing tissues to the limits of their integrity, we carry out a multi-modal study of a simple yet highly dynamic organism, the *Trichoplax Adhaerens*^10–12^, across four orders of magnitude in length (1 *µ*m to 10 mm), and six orders in time (0.1 sec to 10 hours). We report the discovery of abrupt, bulk epithelial tissue fractures (∼10 sec) induced by the organism’s own motility. Coupled with rapid healing (∼10 min), this discovery accounts for dramatic shape change and physiological asexual division in this early-divergent metazoan. We generalize our understanding of this phenomena by codifying it in a heuristic model, highlighting the fundamental questions underlying the debonding/bonding criterion in a soft-active-living material by evoking the concept of an ‘epithelial alloy’. Using a suite of quantitative experimental and numerical techniques, we demonstrate a force-driven ductile to brittle material transition governing the morphodynamics of tissues pushed to the edge of rupture. This work contributes to an important discussion at the core of developmental biology^13–17^, with important applications to an emerging paradigm in materials and tissue engineering^5, 18–20^, wound healing and medicine^8, 21, 22^.

---

Tissues are the paragon example of a ‘smart material’. Cells within a tissue may dynamically reconfigure under stress^15, 23^, exhibit superelastic responses by localizing strain^19^, contract to actively avoid rupture^10^, locally reinforce regions of tissue through recruitment^24^ and other forms of mechanically regulated feedback^25, 26^. Harnessing these properties promises valuable insights for synthetic adaptable materials^18, 20, 27^. Many of these phenomena illustrate the role of mechanical feedback in service of tissues maintaining their integrity under large strains. Here, we address the question – how do cellular tissues behave on the threshold of failure? What determines if a tissue fractures or if it flows? We investigate this question in a ‘minimal tissue system’ that is capable of highly adaptive and fast plastic deformation.

We experimentally study the dynamic epithelial tissues in the marine animal, the *Trichoplax adhaerens*. From the biological perspective, these early-divergent animals have a simple and flat body plan (only 25 *µ*m in thickness but several mm in width), and lack a basement membrane and visible extracellular matrix^10–12^. Their body plan architecture consists of two distinct epithelial tissue layers (top dorsal and bottom ventral, coupled at the edges^10^) which enclose a layer of fiber cells^11^. These animals glide on substrates using ciliary traction^28, 29^. The epithelial tissues consist of millions of cells bound together solely by adherens junctions^30^. From the physical perspective, the *T. adhaerens* is essentially an active composite material composed of flat sheets of morphological distinct epithelial tissues. We leverage the simplicity of these animals and their flat body plan to carry out in-toto imaging experiments that span many orders of magnitude in length and time scales, to gain a fundamental understanding of plastic deformations of epithelial tissue sheets subjected to motility-induced dynamic force landscapes.

Broadly, the plastic response of a wide class of materials can be classified along the spectrum from ductile (ability to draw material into thin wires) to brittle (often hard material likely to break or shatter). There is a rich history of ductile to brittle transitions in non-living matter^31, 32^. The primary mechanisms for tuning through this qualitatively distinct ductile-brittle transition have traditionally been temperature^33^ and disorder^31, 32^. It is an open question whether analogous ductile-to-brittle transitions occur in living tissues and how they may relate to solid to liquid transitions controlled by packing fraction and shape parameters^13, 34^.

In this work, we study entire living animals comprised to two interconnected yet mechanically distinct tissues. In particular, we highlight the contributions of competing forms of connectivity transformations between individual cells and how this competition controls bulk tissue properties such as how and when it yields under load. We complement our experimental efforts by developing a heuristic model designed to focus on the important role of the cell-cell bonding/debonding dynamics in tissues under loading and their dynamic reconfiguration. We derive a simplified model from a powerful existing framework for tissue mechanics^35, 36^ (Methods, Supplementary Information). Our emphasis is on fast time-scale dynamics (∼ sec) in contrast to the more commonly studied case of ∼ min to hour time-scale driving^37^. On these timescales comparable to cell-cell bond lifetime^38^, the dynamics of the network can be represented by a low order model of yielding with dynamic connectivity which we map onto a method in computational geometry (Methods).

Our model reveals a smooth cross-over between ductile and brittle yielding when we vary two key dynamical parameters governing cell-cell bonds: a critical strain for debonding and a time-scale for bond maturation. Controlling a material’s ductile-brittle response to internal/external driving forces is of utmost importance in many engineering applications where catastrophic failure can be more dangerous than gradual yielding in a ductile fashion. However, in some animals such as the *T. adhaerens*, we show that the position in this ductile-brittle spectrum for tissues may be fine-tuned in service of the biological function of plastic shape change and reproduction.

Animals in our lab culture conditions have a wide distribution of amorphous shapes and sizes (Fig. 1A, Methods). Unlike other metazoans, the *T. adhaerens* does not have a fixed stereotypical adult shape. Using long-duration (∼10 hrs) large field-of-view live imaging experiments, we observe that animals undergo time-dependent shape change dynamics from more radially symmetric (circular) shapes to elongated string-like shapes (Fig. 1). These morphological changes are essential for asexual reproduction by fission (Fig. 1B, Supplementary Video 1), which occurs commonly in culture conditions^12^.

**Figure 1.**
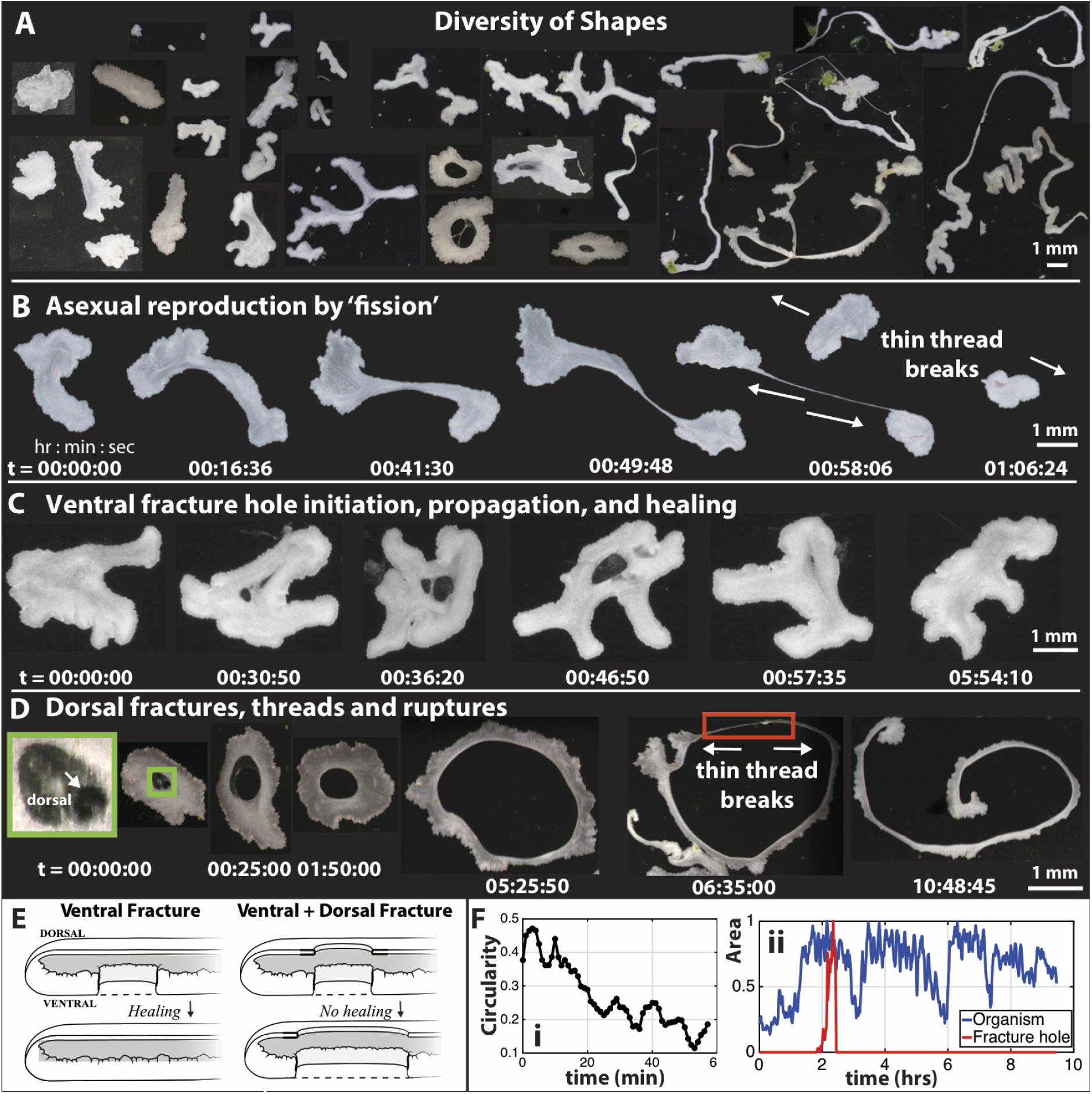
Elastic-plastic shape change phenomena in the *Trichoplax adhaerens*. (A) Lab cultures show a remarkable diversity of amorphous shapes with an extreme variation in size (∼100*µ*m to ∼10 mm). (B) Time-lapse imaging over long time-scales reveal dynamics of this shape change process, eventually leading to asexual fission. The animal forms two regions that pull apart resulting in a plastic deformation (forming a thin thread) which ruptures to give rise to two daughter animals in ∼1 hour (F(i)), reminiscent of ductile necking. (C) In the ventral epithelium, holes appear and disappear (heal) in ∼1 hour (E). Growth via cell-division time-scales are much longer (F(ii)). (D) In addition to ventral holes, we also observe holes in the dorsal epithelium. These dorsal holes do not heal, and lead to through holes inside the animals (E), expanding in area over time. Over time, these torodial animals further break by thread rupture and give rise to long string-like animals.

Surprisingly, in our imaging experiments, we observe holes in both ventral and dorsal epithelial tissues, in native culture conditions. These holes are small in the beginning, grow in size and either heal completely or grow further to the size of an animal (Fig. 1C, D, Supplementary Video 2, 3). Sequential observation of animals reveal a continuous path from flat disc-like sheets to animals with holes (toroidal) which further break into thin, long strings (Fig. 1D, Supplementary Video 3). A contiguous tissue separating into multiple parts requires a topological transformation from a single domain to two or more isolated domains. Even for an elastic sheet this fundamentally requires plastic deformations and neighborhood changes. Hence, we investigate these plastic deformations both at short and long time scales at cellular resolution.

We carried out live cellular scale imaging of the ventral and dorsal tissues in open dish configurations matching native culture conditions (Fig. 2, Supplementary Video 4, Methods). Time lapse imaging over hours reveals that the origins of these plastic events are spontaneous, localized micro-fractures that appear in the ventral tissue (Fig. 2 A,B). These micro-fractures coalesce to form holes with visible cellular debris (Fig. 2B, Extended Data Fig. 2A). In this context of a dissipative tissue, we observe a suppression of fracture propagation with the holes reaching a self-limiting size (Extended Data Fig. 1A). This observation stands out in contrast with classical brittle materials^39, 40^.

**Figure 2.**
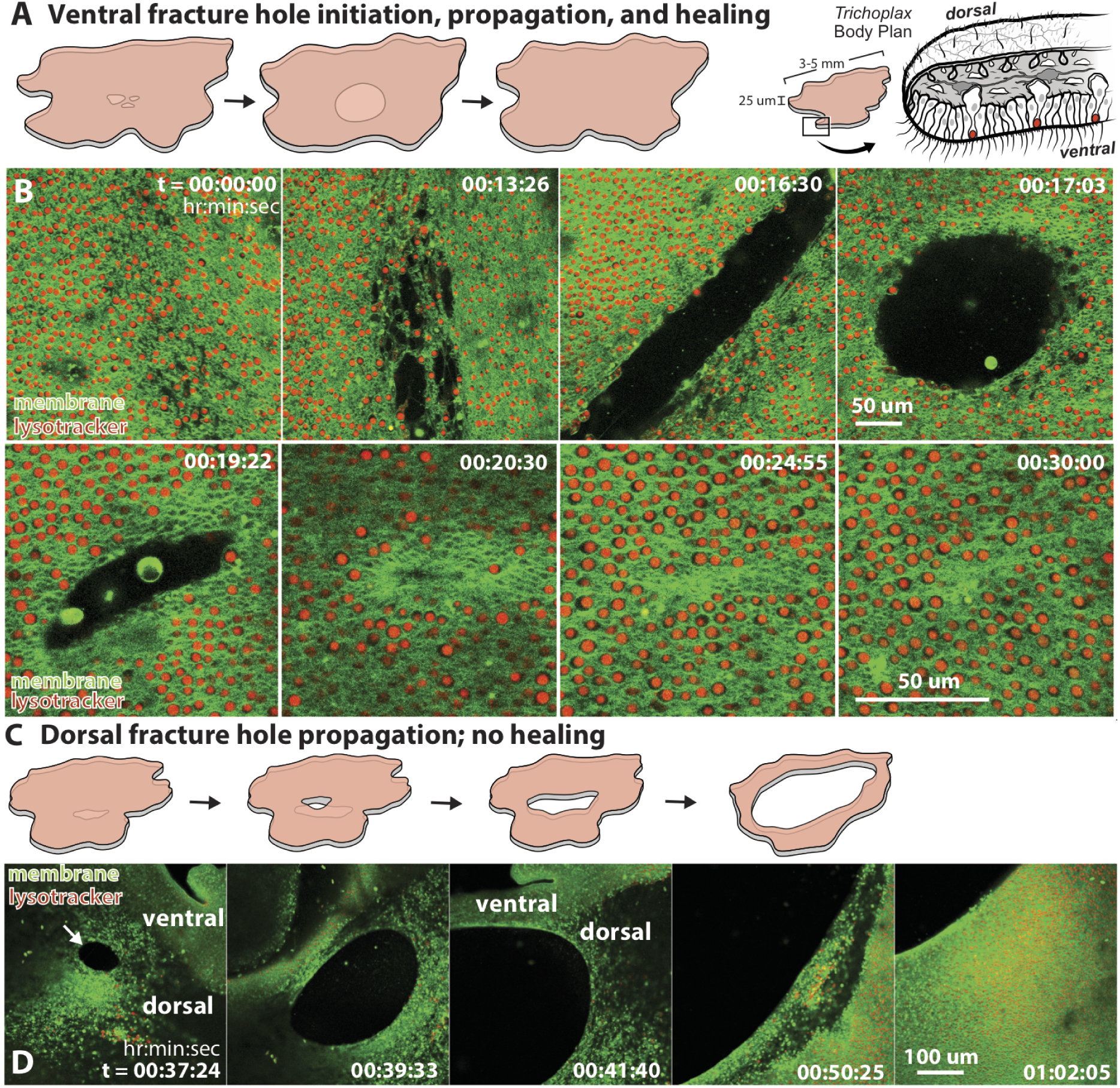
Physiological epithelial tissue fractures in the *T. adhaerens* at a cellular resolution. (A,B) Confocal microscopy reveals that ventral layer fractures initiate as micro-fractures (dark regions in B, N = 19, Methods). These coalesce to form holes with rough edges and broken fragments and further propagate to form a self-limiting larger sized circular hole with smooth boundaries (N = 11). Subsequently, the hole edges come in contact, and the ‘healing’ progresses in a zippering process to completely close the hole. (C,D) Following a ventral fracture, sometimes the dorsal layer can also sustain holes. Dorsal fractures begin as a small, smooth circular hole (white arrow) and expand in size up to the area of the ventral hole below (N = 4). These dorsal holes do not heal and lead to a permanent through hole inside the animal.

Individual cell mechanical heterogeneity can have a strong influence on the collective tissue mechanics^41^. In order to better understand these cellular scale fractures, it is useful to pinpoint the architectural differences between the two major cell types composing ventral tissue: ventral epithelial cells (size ∼ 3*µm*) and the larger lipophilic cells (size ∼ 5*µm*, and comprising 13% of the ventral tissue^42^). Although bi-disperse in size distribution, the ventral tissue does not phase separate but instead remains well mixed. Remarkably, some of these holes can heal completely by a zippering process at the interface of ventral epithelial cells, leaving behind a localized lipophil cell exclusion zone (Fig. 2B, Extended Data Fig. 1C).

This distribution of two cell types in the ventral tissue has a resemblance to traditional alloys in material science^32^. Borrowing from this terminology, we define an ‘epithelial alloy’ as a tissue characterized by multiple populations of mechanically distinct cells (e.g. different size, stiffness, adhesion) in a well-mixed state. In a manner similar to the process of adding carbon into iron to obtain the desired properties^32^, the constitutive behavior of a tissue can be tuned by the percentage of distinct cells present (Fig. 2B, Extended Data Fig. 4).

Focusing on the dorsal epithelium reveals a entirely different phenomenon. Ventral fractures that do not heal can induce a fracture in dorsal layer (Fig. 2C). Unlike in the ventral layer, the dorsal tissue fracture grows quickly into a smooth circular hole similar to the size of ventral hole underneath (Fig. 2D, Extended Data Fig. 1B), akin to a ruptured elastic sheet under tension. Dorsal fractures further lead to the formation of long, narrow threads or string-like phenotypes (Fig. 1D). The interaction between the bulk and peripheral tissue cells can lead to fractures on the edge, and the final breaking of two ultra-thin threads via extension comes down to a single cell-cell junction (Extended Data Fig. 3).

Looking at the dynamics of fracture growth in dorsal vs ventral tissue, we infer distinct mechanical properties under similar strain (Extended Data Fig. 2). This is in line with the fact that dorsal epithelial cells are ultra-thin and flat, while the ventral epithelial cells are smaller and columnar^11^ (Fig. 2A). As ventral and dorsal epithelial tissue layers are coupled via an interconnected fiber cell layer, the picture of a composite tissue emerges. We can consider the properties of these tissue layers to be an integrated composite material, a common framework in engineered materials^43^. The incorporation or layering of units of mechanically distinct materials to constitute a composite can generate highly tunable mechanical properties in living structures^44^. In the context of epithelial tissues, such a composite could permit significant cellular stretch without breaking^19, 45^ or provide rupture resistance complemented by active response to forcing^10^.

Among the existing models to study tissue mechanics, cell-resolved tissue models have become a powerful tool for understanding rigidity transitions^41, 46, 47^, active tension networks^24^, the importance of apical-basal symmetry breaking at the cellular level^48^, how embryos generate shape^36^, and the interplay of fluidity and healing of ablated wounds^49^. Broadly, the majority of existing models have been developed to study tissue response to slow forcing timescales in confluent mono-layers with a single cell type. Our experimental observations suggest that there is more to learn about tissue mechanics by studying tissue response at unsteady and fast loading timescales (comparable to the cell-cell junction lifetime), without a confluence constraint in multi-component cellular tissues.

Even the simplest living tissue represents a complicated interplay between cell shape, compliance, flow/remodeling, adhesion, heterogeneity, and even mechanobiological feedback. When faced with such complicated construction, one approach for progressing seeks to extract an emergent simplicity. This ambition often leverages a few primary methods: separation of timescales, dominant effects, and low-order expansions. The thoughtful application of these methods has generated many useful models which have enriched our understanding in many interesting regimes of tissue behavior^35, 50^. Detailed experiments/numerics of soft-sticky granular matter suggest that system wide strain arises not from the strain of the individual elements but the rearrangement of the network in regions called Shear Transformation Zones (STZs)^51–53^. Yet, experimental studies on epithelial tissues suggest that cell strain can account for tissue strain^3, 54, 55^ and that yielding comes from higher order cellular mechanics such as cytoskeletal relaxation and cortical depletion^19^. This conundrum lies at the very heart of how we think about tissues under load: on short timescales, do tissues yield by fracture or by flow?

In an effort to distill this essential discussion, we reformulated this central question as a competition between edge-formation and neighbor exchange mediated by cell-cell junction forming kinetics in a disordered model of an active tissue inspired by adhesive granular matter. In order to ground our model in experiments, we identify some key observable features that characterize epithelial tissues in the *T. adhaerens*: these include cell size, cell spatial distribution, distributed traction mediated propulsion^28, 29^ and non-confluence. Our experimental observations encouraged us to reconsider the often useful assumptions: (i) local cell mechanical uniformity, (ii) tissue confluence, and (iii) the surface energy of cell-cell interactions constructed from a seperation of timescales (Extended Data Fig. 4, Methods).

We capture fracture and healing phenomena by modeling the bonding and debonding of cell-cell junctions. A good starting point for compliant adhesive cells is the vertex energy function^50^, where we unfold the surface tension-like term into the contribution coming from cortical tension and the contribution coming from interaction with other sticky cells^56, 57^. Since the dynamics we study overlap with the timescales of the cell-cell bond lifetime, we split the second term of cell-cell interaction energy into the dynamics of adhesive bonds. The energy function for a unit cell *i* is given by:

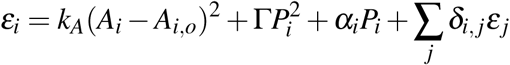

Where *k*_*A*_ and G are the areal (A) and perimeter (P) rigidity respectively, *α* is the cell’s cortical tension, *δ*_*i, j*_ is a dynamical term counting the number of cell-cell junctions present and bound between cell’s *i* and *j* where we sum over all the cells in contact with cell *i*. Our proposed model replaces the complexities of cell shape with an effective compliance to clarify the central role of the competition between edge-forming and STZ transformations. The mechanical properties of this model tissue will be determined by the cellular compliance and the bonding and debonding time scales (Methods, Extended Data Fig. 4).

A classical model for the force dependent unbinding of two state systems in a tissue context follows Bell’s seminal work^58^. When we study an ensemble of these two-state junctions, we find an critical behavior where the release of one cell-cell junction redistributes that load to the others, slightly increasing the remaining junction’s probability of transitioning. This induces a cascade of events which ruptures the cell-cell junctions abruptly. We find that for sufficiently high affinities and ensemble sizes, the collective dynamics is well approximated by a Heaviside-step (Supplementary Information):

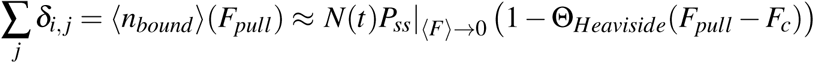

Where ⟨*n*_*bound*_⟩ is the average number of bound junctions, *F*_*pull*_ is the average pull force, *N*(*t*) is the number of local Cadherins clustering^59, 60^ (timescale ∼ 10 sec), and *F*_*c*_ is the critical failure force. *N*(*t*) is the limiting timescale of our proposed bonding criterion. We define this timescale as the maturation time *τ*_*mature*_ of the cell-cell junction^61^. In the context of an elastic material, the force threshold can be translated to a geometric threshold. Approximating the cell compliance as elastic is reasonable on sufficiently short timescales^25^.

Next, we focus our study on the cell-cell junction dynamics instead of the cells themselves by replacing our vertex degrees of freedom with a simple (two-body) Hertzian cell compliance under compression and harmonic response under tension. This allows us to map our model onto the dynamics of the connectivity matrix emphasizing the debonding/bonding criteria over the cell shape dynamics. This model can be interpreted as a two-dimensional, bi-disperse, adhesive granular matter model with slow (comparable to the timescale of the dynamics) bond maturation kinetics^62^. New bonds begin their maturation when they are neighbors in a Delauney Triangulation and are within the geometric threshold, taking the form of a finite distance Delauney network (Extended Data Fig. 4, Methods, Supplementary Information).

With this heuristic framework in place, we can begin subjecting the model tissue to a controlled distributed actuation^28, 29^ (Methods). We conduct two primary studies of the response of this network. First, we study the model under fixed stretching for a finite time (Fig. 3A). We vary the threshold strain for breaking the cell-cell bonds and sweep through a range of driving force gradients. We observe three characteristic responses. The first is the presence of a yield stress, where below a given driving amplitude, the material response is purely elastic, while above a certain threshold, plastic deformations are observed (Fig. 3A, Supplementary Video 5). Further, we compare our adhesive granular material model to the classical phenomenological Herschel-Buckley model for a yield stress fluid^52^, and find a good fit with a power-law scaling of ∼ 3 (Fig. 3B) (Methods).

**Figure 3.**
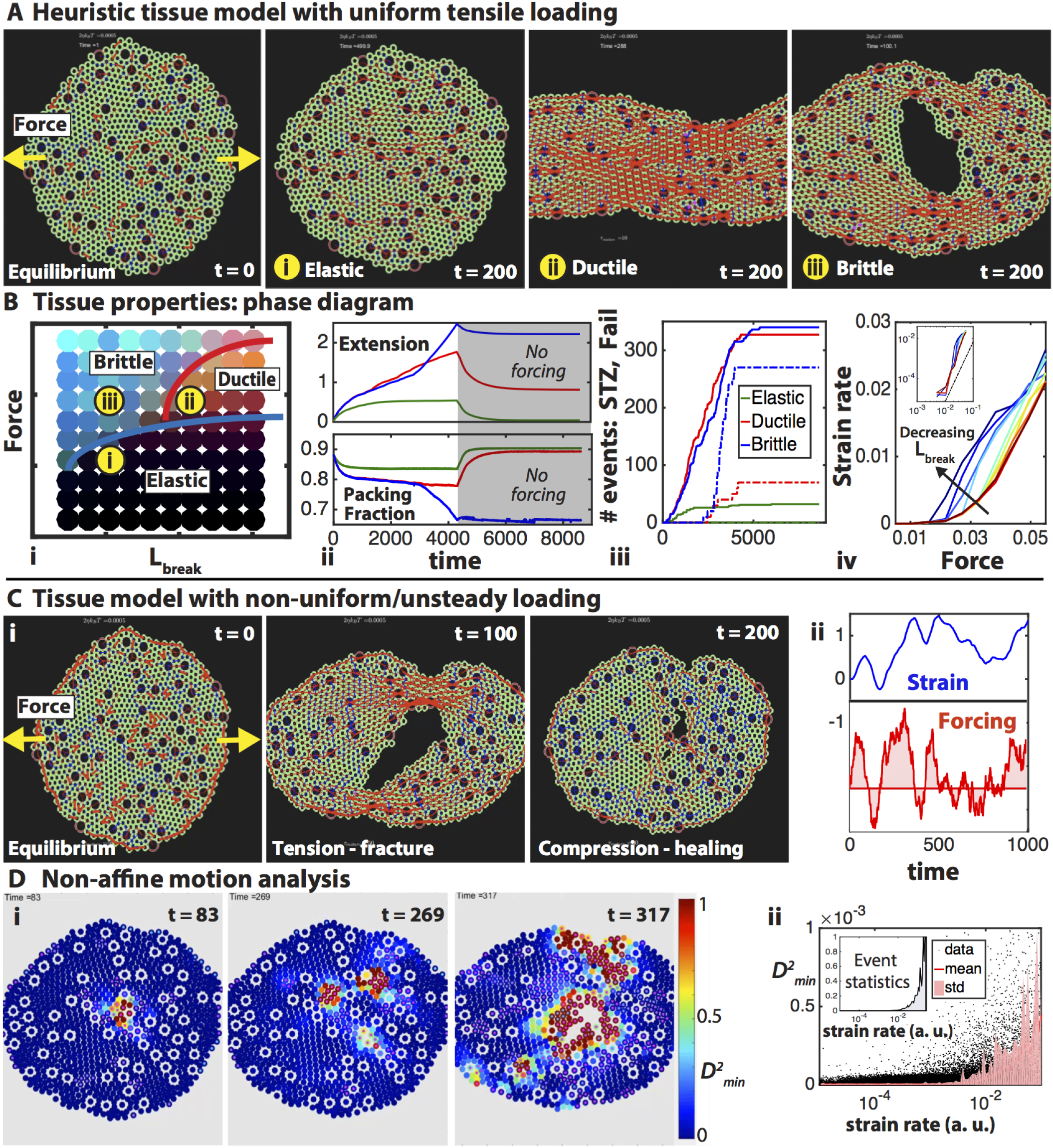
A heuristic in-silico tissue model captures yielding and ductile–brittle transitions. The model consists of soft balls (cells) connected via dynamic, compliant junctions (adhesion) (Methods, Extended Data Fig. 4, Supplementary Information). (A) The system is driven out of equilibrium by a uniform tensile force gradient to mimic the lowest-order devioric component of distributed propulsion (yellow arrows). At low force gradients, we observe elastic behavior A(i). At high force gradients, we observe two qualitatively distinct plastic regimes – ductile regime A(ii) and brittle regime A(iii) with tissue fractures (Methods). (B) (i) We run a parameter sweep by varying driving force gradient, 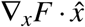, and the strain threshold for breaking the cell-cell junctions, *L*_*break*_, to generate a phase diagram that clearly delineates three regimes of tissue properties: elastic–ductile–brittle (Extended Data Fig. 5). (ii) At the whole tissue scale, we characterize the response with the extension and packing fraction versus time. (iii) At the cell-scale, we monitor two classes of transformations to the cell-cell connectivity associated with edge-forming transformations (dotted) and constraint-count-conserving transformations (solid) akin to STZs (Methods). (iv) The steady state response of each parameter set is consistent with the Herschel-Buckley response with a finite yield stress and a scaling of 3. (C) (i) To investigate the role of dynamic loading, the model is driven by non-uniform, unsteady, fluctuating forces with a characteristic timescale (ii) reminiscent of locomotion in the *T. adhaerens* (Methods). This time-varying force landscape captures both tissue fracture and healing dynamics. (D) (i) To identify experimentally accessible measurements of yielding, we carry out a non-affine motion analysis using the metric 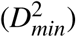 ^53^ on the simulations (Methods). (ii) The probability of high non-affine motion 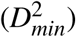 plotted against the instantaneous strain rate highlights signatures consistent with a yield stress behavior.

In order to classify the yielding behavior of this model tissue, we can define a ‘brittle’ material as one which undergoes transformations that create new edges without STZs. A ductile response can still fracture but will do so with STZs as precursors to rupture. We use this to develop a technically precise definition to characterize our ductile-brittle spectrum: constraint-count-conserving (CCC) transformations compare the number of constraints before and after a cell-cell neighborhood transformation. If the number of constraints is locally conserved we call it a CCC transformation.

In the second regime, above the yield transition, we observe the tissue going through many CCC transformations in the neighborhood matrix. These CCC transformations are analogous (though not precisely equivalent to) to T1 transformations within a tissue. This regime corresponds to our more ductile-like yielding.

The third outcome of pulling on this model tissue is producing a large population of non-CCC transformations indicating the generation of new boundaries. This type of yielding can be understood as a more brittle fracture via loss of tissue continuity and few associated CCC transformations to relax the energy (Methods, Supplementary Video 5, Methods). By consolidating these results grouped by the distributed force gradient versus debonding criteria, we draw a phase diagram where the material undergoes ductile to brittle transition above its yield stress (Fig. 3B, Extended Data Fig. 5, Supplementary Video 5, Methods).

We propose thinking of these phenomena as a heuristic analogy to a ductile to brittle transition within a toy model of cellular tissue. A brittle material yields predominately by fracture (new edge formation) and a ductile material yields by flow (or STZs). The qualitative ramifications suggest that it may be biologically feasible to tune heuristic parameters in such a way to make a specific tissue ductile (such as dorsal epithelium) while another tissue to be more brittle (such as ventral epithelium) under the same driving force. A tantalizing outcome is that the composite tissues of the *T. Adhaerens* may be qualitatively understood by learning from the continuous crossover in this simple model.

Observing these animals under a microscope clearly reveals an unsteady nature of the driving force induced by motility (Extended Data Fig. 8). Next we drive this model tissue under a long wavelength collective mode akin to the unsteady dynamics observed which captures the self-inflicted rheology of the tissue. We implement this by varying the pulling amplitude in time using a 1D random walk in a soft harmonic potential to capture both the characteristic timescales and stochasticity of the driving. Under such unsteady loading, the model tissue deforms to capture both fractures and healing (Fig. 3C, Supplementary Video 6, Methods).

The ambition of the heuristic model is to refine our questions and enrich experimental quantification. We can leverage the omniscience of the model to quantify changes to the connectivity matrix of the cells to arrive at experimentally observable parameters. This capability allows us to identify relationships between connectivity changing events and locally non-affine motion. Existing metrics for quantification of non-affine motion such as 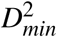 ^53^ (Extended Data Fig. 6, Methods) excel at capturing the localization of Shear Transformation Zones (STZ) in space and time. These signatures form a compelling measurement of dynamical heterogeneity^63^, and we exploit these signatures to provide support for a finite yield stress within the tissue. We approach this challenge by first mapping the presence of large 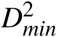 values onto the connectivity changing events. We find that 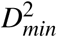 is an excellent indicator of these events (both CCC and non-CCC) (Fig. 3D). Next, we find that the yield stress character of this tissue manifests itself as an abrupt rise in the correlational measure between 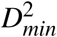 (measure of STZ) and the instantaneous strain rate (Fig.3D).

Inspired by results from the heuristic in-silico tissue model, we turn back to experiments to further understand how forces govern tissue mechanics in the *T. adhaerens*. In order to perform quantitative measurements in live epithelial tissues in native conditions, we developed novel cell tagging techniques and computational data analysis techniques (Extended Data Fig. 7, Methods). As suggested by our model, the experiments indeed confirm that organism-scale motile forces can give rise to brittle-like tissue fractures (Fig. 4).

**Figure 4.**
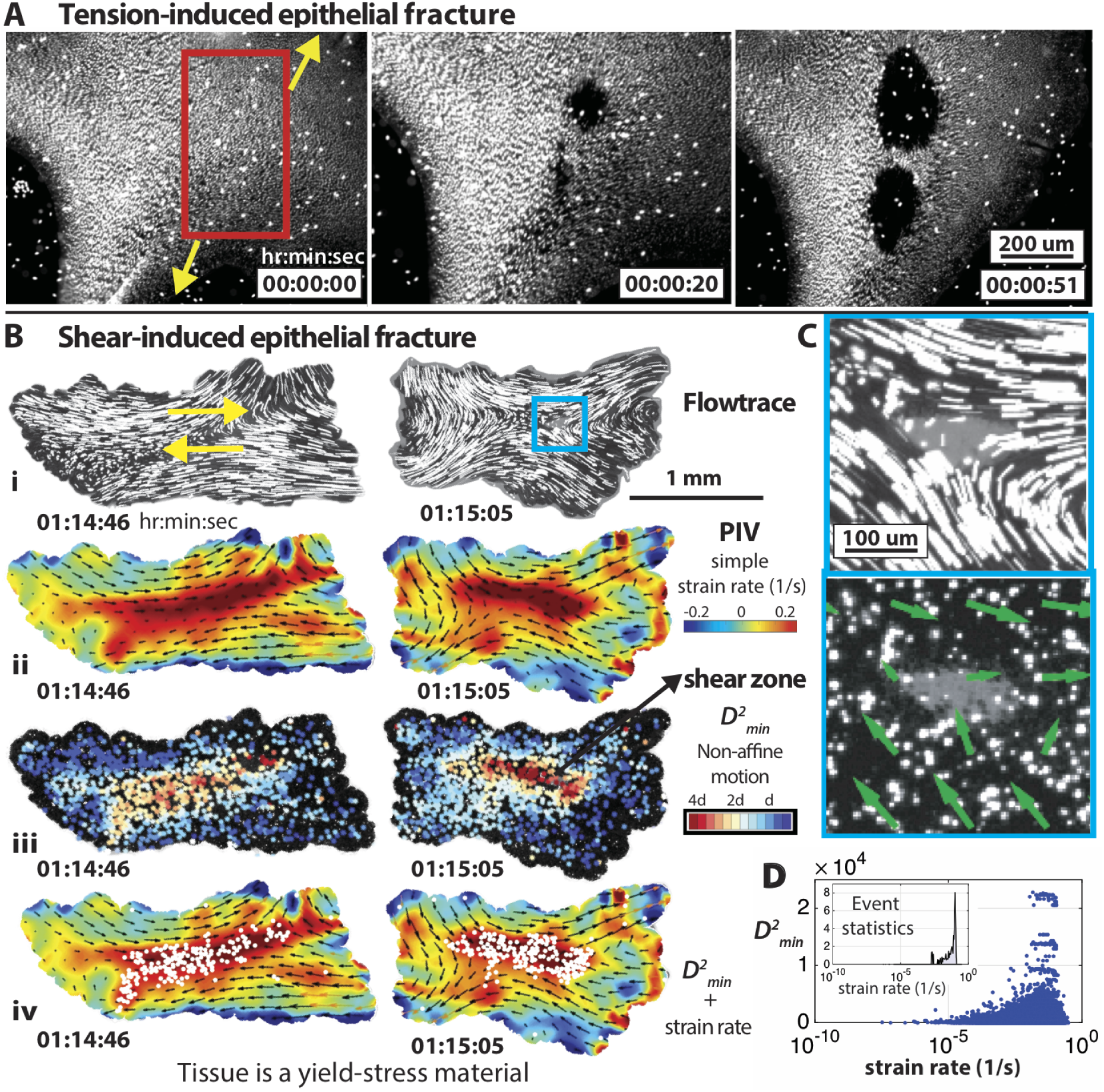
Organismal motility-induced forces cause local tissue fractures in the *T. adhaerens*. (A) Fluorescence microscopy reveals that Tensile forces can induce fast (∼2 min) brittle-like fracture dynamics in the ventral epithelium (Extended Data Fig. 9, Methods). Micro-fractures rapidly coalesce to form larger holes. (B) While simultaneously imaging sticky fluorescent micro-beads on the dorsal epithelium, we observe a shear-induced ventral fracture (Extended Data Fig. 7, Methods). (i) Flowtrace^65^ visualizations reveal bead trajectories in a region of local shear-induced fracture (zoom-in insets (C)). (ii) The internal strain rate (with peak ∼0.2 *s*^*-*1^) contours from a PIV analysis overlaid on the velocity vectors. (iii) Non-affine motion analysis metric 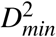 ^53^ captures high shear transformations (Methods). (iv) Regions of high 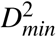 (white dots) show excellent correlation with regions of high internal strain (red contours). (D) Quantification of 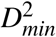 versus the strain rate, demonstrates signatures of a yield-stress material – consistent with model predictions (Fig. 3D).

The principal modes by which materials fail by fracture are classified as tensile failure (mode I), shear failure (mode II) and failure due to out of plane tear (mode III)^40^. We find experimental evidence of the first two failure modes in the tissues of *T. adhaerens*: a tension-induced failure (mode I) (Fig. 4A, Extended Data Fig. 9, Supplementary Video 7, Methods) and a shear-induced failure (mode II) (Fig. 4B,C, Supplementary Video 8, Methods). In our experiments, we do not observe the third failure mode (mode III) since the forcing is confined to a two-dimensional plane.

Next, we proceed to investigate the material properties of these tissues. In order to do this, we first experimentally measure the amount of plastic deformation (shear transformations) in the tissues using the non-affine deformation metric 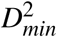 ^53^ (Fig. 4B, Supplementary Video 9, Methods), encouraged by its success in our model above. Secondly, we measure the internal strain rate (typically 20% *s*^*−*1^; maximal rupture strain ∼ 300%) induced in the tissues due to the animal’s own motility (Extended Data Fig. 8, Methods). We find a high degree of correlation between these measures of plastic deformation and internal strain rate (Fig. 4B, Supplementary Video 10). In both our experiments (Fig. 4D) and model (Fig.3D), we observe an abrupt rise in the 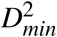 versus strain rate plot, which strongly suggests that these epithelial tissues exhibit properties of yield stress materials both with and without new-edge formation. Our measurements compare well with other yield stress phenomena in biological tissues, including zebrafish embryos^13^ (∼ 75% linear strain) or in cell culture systems^19^ (∼ 300% areal strain).

We have so far observed that the *T. adhaerens* experiences loading at fast time-scales. Our model reveals that under such fast loading time-scales (relative to cell-cell maturation time *τ*_*mature*_), otherwise ductile tissues can become brittle since new cell-cell bonds are still fragile (Extended Data Fig. 6). It is therefore noteworthy that a tissue under such fast loading can yield by flow, in contrast to yielding by fracture. This suggests that *T. adhaerens* might have tuned its tissue properties via making epithelial alloys and composite architecture. Our observations of the ductile characteristics of the dorsal layer and brittle-like properties of the ventral layer further support this idea.

The rich and diverse dynamics of a living tissue can be interpreted through many complementary viewpoints – each with their own useful insights. Here we have suggested a perspective combining adhesive granular matter and concepts from materials science to study the fast timescale dynamics of epithelial tissues under distributed loading. We report the utility of this approach to inform the generic aspects of the yielding behavior of cellularized tissues, and the competition between fracture and flow. This perspective complements a broader trend in the biological physics of tissues^13, 19, 22, 24, 35, 37, 57^.

Living materials such as epithelial alloys arise from repeated evolutionary trial and error. On evolutionary timescales, epithelial alloys may allow for tuning of the emergent mechanics of tissues to avoid catastrophic failure. Here we have observed how tissues can localize damage by increasing dissipation and by the ability to rapidly heal. This resilience to failure modes can expand the ecological niches where organisms can thrive, enabling the early divergent *T. adhaerens* to successfully populate tropical oceans worldwide^64^.

## Acknowledgements

We thank A. Bhargava and N. Eiseman for their contributions in the initial stages of the project, and S. Armon, P. Vyas and members of the Prakash Lab at Stanford University for useful suggestions. We thank V. Chikkadi, S. Bandyopadhyay, S. Shekhar, C. Lowe for helpful scientific discussions; K. Uhlinger for help with marine algae cultures; L. Buss for the generous gift of animals; R. Konte for help with the illustrations, C. Espenel, A. Khan, and J. Mulholland (CSIF, Stanford University) for help with microscopy. M.S.B. was supported by the National Science Foundation Graduate Research Fellowship (DGE-1147470) and the Stanford University BioX Fellows Program. This work was supported by NSF CCC grant (DBI-1548297) (M.P.), CZI BioHub Investigator Program (M.P.), the Howard Hughes Medical Institute (M.P.).

## Author contributions statement

All authors designed the research; V.N.P. performed the experiments with assistance from M.S.B. and M.P.; V.N.P. and M.S.B. analyzed the data; M.S.B. developed the theoretical framework and performed simulations; All authors wrote and reviewed the manuscript.

## Methods

### Laboratory cultures of *T. adhaerens* and marine algae

We maintain lab cultures of *T. adhaerens* (gift from Leo Buss, Yale University) in shelves of Petri dishes containing artificial seawater (ASW) in a dedicated room maintained at 19°C temperature with an 18-hour light and 6-hour dark cycle. The ASW is prepared by adding measured quantities of Kent Marine Reef Salt mix in Millipore water on a magnetic stirrer in order to reach a salinity of 3%. The salinity is measured using a handheld salinity probe (CDH45, Omega Engineering). The food source for the *T. adhaerens* is the marine algae *Rhodomonas lens*. We maintain separate axenic cultures of R-lens in ASW inside 500 ml Erlenmeyer flasks in dedicated incubators maintained at 14°C temperature. The algae are provided with nutrients (Micro Algae Grow, Florida Aqua Farms) and an 18-hour light and 6-hour dark cycle for optimum growth. We carry out regular maintenance of the animal and algae cultures once every week. We prepare fresh plates by filling three-fourths the volume of a petri dish with fresh ASW and algae nutrients (250 *µ*l nutrient per liter of ASW). We then seed these fresh plates with about 5-10 ml of R-lens algae and let the plates grow an algae biofilm for 1-2 days. After this, we transfer about 20-30 animals from another slightly older dish and let the animals settle down and grow. The ASW in older plates of T. adhaerens are replaced every week (2/3 volume) with fresh ASW, algae nutrients and a few ml of R-lens algae. The axenic algae cultures are split once every week; 50 ml of algae from an older flask is transferred into 200 ml of fresh ASW and nutrients (with concentration of 500 *µ*l nutrients per liter of ASW). These axenic algae culture flasks are shaken regularly at least once every day.

### Experiments: Live Imaging and Microscopy

For long-term imaging studies (Fig. 1), the lab culture petri dishes containing *T. adhaerens* are placed on a base of a modified Stereomicroscope (Leica S9 D) stand; where a digital SLR camera (Canon EOS Rebel T3i) fitted with a zoom lens (Canon EF lens 24-105 mm) replaces the stereo-zoom optical magnification unit. The imaging system employs a quasi-dark-field setting and high-resolution images are acquired remotely on a computer once every 5 seconds.

The animals in culture dishes adhere to the bottom surface, and we gently impinge them with small water jets from the side repeatedly using micropipettes in order to peel them off the surface and transfer them elsewhere for imaging. Depending on the type of experiment, we carry out live imaging of the animals by transferring them into different imaging chambers. We employed chambered coverglass wells: Nunc Lab-Tek chambers (1-well, Thermo Fisher Scientific) – this configuration allows imaging of the animal under natural conditions without any confinement. We then stain the animals with fluorescent labels and carried out live confocal imaging (Zeiss LSM 780 and 880) for many hours, using minimum laser intensity (1-5%) and an acquisition rate of 2 fps, with some variation in acquisition rate depending on objective used (10X air, 20X air, 25X water, 40X oil).

We also carried out imaging in custom-designed PDMS milli-fluidic hexagonal chambers (thickness 40 *µ*m, and widths ranging from 1-3 mm); these chambers confine the animal movements in z-direction, and allow motility only in the two dimensional xy plane, depending on the width of the chambers. Once the animals have settled down in the imaging chambers (over hours) and the chambers are sealed, they are mounted on an inverted microscope (Nikon TE2000-U) and imaged under bright field and fluorescence (lambda XL light source) modalities. The images are captured using on a high-speed camera (ORCA Flash 4.0, Hamamatsu) generally at frame rates of 10 fps for long durations (many hours), but the frame rates and durations are varied in some specific experiments. Depending on the particular experiment’s objective to achieve specific fields-of-view, different objectives – 2X, 4X, 10X and 60X oil immersion objectives were used for imaging.

### Numerical study of heuristic model

Given the complexity in tissue mechanics, the best choice of model is one which focuses our attention on the new representation: cell-cell debonding/rebonding criteria. The simplest model which captures these phenomena can be effectively characterized as a ‘sticky-ball’ model. Cells are represented by soft circles that can deform under a Hertzian potential (Extended Data Fig. 4). Cells are then linked through geometrically defined dynamical network of harmonic springs when under extension. The neighborhood state governing the kinetics of the transition between neighborhood configurations is seeded by the finite length Delauney transformation. Springs capture the lowest order nonlinearity through a yield threshold

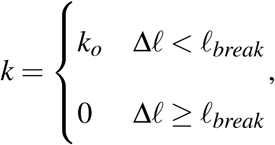

withΔ*l* = |*r*_*i*_ − *r*_*j*_|.

This is a representation of maximal strain yielding (Supplementary Information). In the case of harmonic springs, this is simultaneously consistent with the maximal stress and Von mises criterion through the devioric moments relationship to energy density. Two populations of cells of differing sizes are initialized to suppress crystallization (consistent with experimental observations). To ensure that the cell populations are uniformly distributed throughout the tissue, we leverage the Differential Adhesion Hypothesis (DAH)^56^ encoded through attachment stiffness between cell types. By choosing the correct relationship, *k*_22_ < *k*_11_ < *k*_12_, cells will preferentially distribute throughout the tissue when rapidly quenched from a random initial state (but not an infinite–temperature quench). The quench dynamics take the form of a gradient descent on the energy landscape with the dynamical equation:

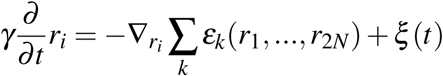

The gradients are taken analytically and the positions are updated using a combination of Euler’s method and a custom variant of a fast inertial relaxation engine.

The number of metastable states in the model tissues explodes rapidly with increased bidispersity via size and percentage of large cells, reminiscent of a collodial glass. In an ensemble of these metastable states, we can study the packing fraction, the hexatic order and the residual force as a function of the size difference between cells and the percentage of each cell population. We set these to be consistent with experimental measurements and observe similar observables including the orientational correlation length of the hexatic order parameter (Extended Data Fig. 4). Even though our model does not have viscoelastic components, we find emergent bulk viscoelasticity at finite temperature through neighborhood exchanges consistent with many yield stress fluids^52^.

A classic phenomenological model for a yield stress fluid is captured by the Herschel-Buckley equation 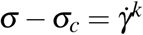(where the steady state flow rate is equal to the kth-root of the stress above the yield stress)^66^. We can compare our adhesive granular material model to this classical behavior in the long-time limit by fitting the strain versus time to a curve (with fit parameters *A, B* and *τ*_*relax*_ = 45 simulation units) of the form: *δ* (*t*) = *A* + *Bt*(1 *- e*^*-t/τ*^*relax*) (Extended Data Fig. 5). By plotting the asymptote of the fit versus the fixed driving force, we find that the Herschel-Buckley equation is consistent with a *k* ∼ 3 scaling (Fig. 3B(iv)).

To mimic the distributed activity of the organism induced by motility, we expand the distributed locomotion to lowest order in the devioric field^28^. These long wavelength collective modes represent a low order approximation to the dynamics of the forcing on this epithelial layer arising from distributed propulsion. This has the added benefit of serving as a control parameter for our driving away from equilibrium. By disentangling the dynamics from the distributed forcing, we can more easily study the response of the model tissue to steady state forcing. The functional form for this forcing takes the form of:

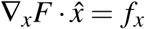

Where we use *f*_*x*_ to characterize the amplitude of the gradient in the x-component of activity. The study of the model under applied distributed propulsion coefficient *f*_*x*_, begins with a constant pull *f*_*x*_ followed by activity *f*_*x*_ = 0 after *t* = *T*_*stop*_. The response of the model is characterized as a function of *f*_*x*_. We have also carried out complementary studies using a shear amplitude which qualitatively showed a similar behavior (data not shown).

To classify the response of this material as a function of amplitude *f*_*x*_, *τ*_*mature*_, and *ℓ*_*break*_, we developed a suite of microstate measurements to characterize the type of transformations incurred by driving it out of equilibrium. The first two measures are the types of transformations to the neighborhood matrix. In a STZ, a constraint broken is replaced with another locally. The local number of constraints does not change with an STZ. In contrast, a new edge forming transformation reduces the number of constraints and picks up the modes associated with a new edge. By counting the number of constraints locally, we can classify changes to the connectivity as either constraint-count-conserving (CCC) or non-CCC. These technical definitions open the door to the characterization of the competition between fracture and flow.

In addition, we define the packing fraction using a Monte Carlo method for each configuraiton. We place 10^6^ points randomly within the domain of the material asking if each is within a cell or within the boundary of the organism. The number which land inside a cell over cardinality of the set landing in the cells unioned with those in the organism provides a calculation of the Packing Fraction. The parameter spaces (Fig. 3B, Extended data fig. 5) were generated by superimposing these three measures. Taking the practical definition of a ‘brittle material’ as one which overwhelmingly creates new edges and the ‘ductile’ material overwhelmingly changes neighbors, we can characterize a crossover from ductile-like materials at high *ℓ* _*break*_ (short *τ*_*mature*_) to more brittle materials at low *ℓ* _*break*_ (long *τ*_*mature*_).

To study the role of the dynamic force landscape on the evolution of the connectivity, we made our amplitude dependent upon time, *f*_*x*_(*t*). The dynamics of this driving amplitude took the form of stochastic Langevin dynamics governed by the equation:

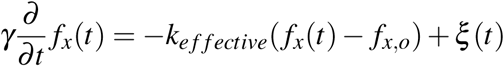

With *ξ* (*t*) = 2*kTγδ* (*t –t*^′^) consistent with fluctuation-dissipation theorem. This mimics an overdamped particle in a harmonic well, with a characteristic timescale of response determined by the relaxation time of the system.

By driving our model tissue with this stochastic distributed driving, we can measure the accumulated morphodynamics attributed to this dynamic force landscape. We observe three central phenomena: fracture, STZs, and healing. Since we want to port our understanding to experiments where we do not have direct and error-free access to the microstate, we sought to characterize the dynamics of transformation in a coarse grained way using a metric from the granular matter literature, 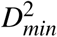. By comparing the signatures of high non-affine motion to known neighborhood transformations, we extracted a threshold above which 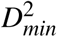 is a good indicator of neighborhood transformation (Fig. 3). We also used the model dynamics to characterize the sensitivity of these measurements to parameter selection confirming best practices of choosing the interrogation radius consistent with the point correlation function (Extended Data Fig. 6).

### Experiments: Tagging Techniques

We use a fluorescent LysoTracker dye (deep red, Thermo Fisher Scientific) in 1:500 concentration in ASW to stain granules (acidic inclusions) in lipophil cells of the ventral epithelium^11^ in the *T. adhaerens*. The cell membrane is stained using the fluorescent FM 1-43 dye (Thermo Fisher Scientific) in 1:100 concentration. We developed an assay to tag the dorsal epithelium using sticky fluorescent micro-beads (micro-spheres). Fluorescent carboxylate-modified micro-spheres (0.5 *µ*m Red, Thermo Fisher Scientific) are coated with a protein – Wheat Germ Agglutinin, WGA (Alexa Fluor 350 conjugate, Thermo Fisher Scientific), which is a lectin that specifically binds to cell membranes. In this step, we used a protein coupling kit (Polysciences) to coat micro-spheres with WGA^14^. This sticky microbeads assay involves an incubation step that lasts overnight (10-12 hours) and in the meanwhile, we prepare the animals for imaging experiments. The animals that are selected for imaging (∼5) from the culture petri dishes were transferred into a fresh dish with only ASW. We ‘starve’ these animals in ASW for periods of 4-6 hours without providing algae – during this period they unfold and settle down on the glass substrate and start gliding around to explore the environment. In this starvation period, the animals shed most of their algae biomass ‘cargo’ that they naturally accumulate in culture conditions. This step greatly enhances our ability to image their epithelial tissue layers with clarity. To ensure their health, these starved animals are now fed small amounts of R-lens algae (few ml) for about an hour before sprinkling the sticky beads.

After preparing the sticky microbeads, we sonicate the samples for ∼5 minutes to break down aggregates, followed by dilution (1:10) in ASW. We now sprinkle this dilute solution of sticky beads on the animals by locally (near their dorsal epithelium) delivering 10 *µ*l volumes repeatedly (∼5 times) using a micropipette, in intervals of 30 minutes over a period of 2.5 hours. The animals are ‘washed’ twice in order to ensure that we retain only the beads that permanently stick on them for our imaging. The washing process involves transferring the animals into fresh ASW dishes and letting them settle there for few hours. Once again, the animals shed all extraneous materials (such as bead aggregates) during these starvation periods. After the washing process, the animals are then transferred onto the PDMS chips in a droplet, and once they unfold and settle, a cover slip is gently placed on the chip and sealed in order to minimize evaporation, and mounted on the microscope for imaging.

### Computational Analysis Techniques

We used different computational analysis techniques to quantify the experimental time-lapse datasets. The dorsal layer datasets with sticky-microbeads tagging were visualized using Flowtrace^65^, a simple tool for visualizing coherent structures in biological fluid flows (in ImageJ). The Flowtrace algorithm generates pathlines of particles with a given input projection time (here we chose 5 seconds).

The dorsal layer sticky-beads datasets were also analyzed using the Particle Tracking technique in MATLAB (Mathworks). The first step involves image-processing operations such as background subtraction and spatial filtering to ensure that we can optimally identify the locations of the fluorescent ‘blobs’ of sticky-microbeads for tracking. In the next step, the particles are tracked over time using the nearest-neighbor particle-tracking algorithm. The particle tracks are then stored for further data analysis.

Next, we studied the non-affine motion (non-uniform or disordered motion) of these tracked micro-beads to measure the local amount of deviation in particle displacements from a linear strain field. Non-affine particle motion is a key feature in many soft matter systems such as amorphous solids^53^, colloids^67^, jammed materials^68^, granular materials^69^ and cell migration^70^. We quantified non-affine motion using the 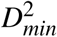 metric^53^ for all the tracked particles over a short time-scale of 1 second. This 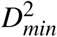 metric^53^ minimizes the mean-squared difference between the actual displacements of the neighboring particles relative to a central one and the relative displacements that they would have if they were in a region of uniform strain:

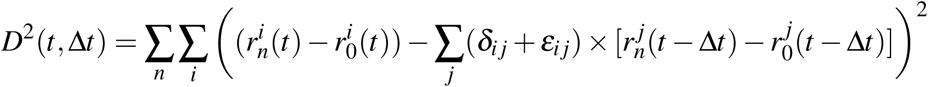

where, 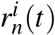 is the *i*th component of the position of *n*th particles at time *t*. The uniform region of strain *ε*_*i j*_ that minimizes *D*^2^ is then calculated according to Falk & Langer^53^. Then 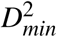 is the minimum value of *D*^2^(*t*, Δ*t*), which is the local deviation from affine deformation during the time interval [*t -* Δ*t, t*]. The output of this metric is a squared length-scale that quantifies the amount of disorder in the particle motion. The results are sensitive to the selected size of radius around the particles, and we have chosen optimal values of this parameter for our data analysis.

We employed Particle Image Velocimetry (PIV) analysis (PIVlab package^71^ in MATLAB) to quantify the flow-fields in the dorsal layer sticky-microbeads time-lapse datasets in large width (13×13mm square) confined PDMS milli-fluidic chips with variable fields-of-view. The animal is tracked manually for fixed durations, and the time-lapse recording is stopped when the field-of-view is changed. The PIV analysis is carried out for specific duration segments of these datasets where the field-of-view is fixed, and over short time-scales (1 second). The image preprocessing step involves high-pass filtering, and we then carry out the PIV analysis by dividing the images into 64×64 pixel interrogation windows with 50% overlap on the first pass, and we use 32×32 pixel interrogation windows with 50% overlap for the second pass. Next, we select velocity threshold limits for post-processing the resulting velocity vector fields, smooth data and use interpolation for missing vectors. We then calibrate the results, and calculate derived quantities such as the simple strain rate.

---

**Extended Data Figure 1:**
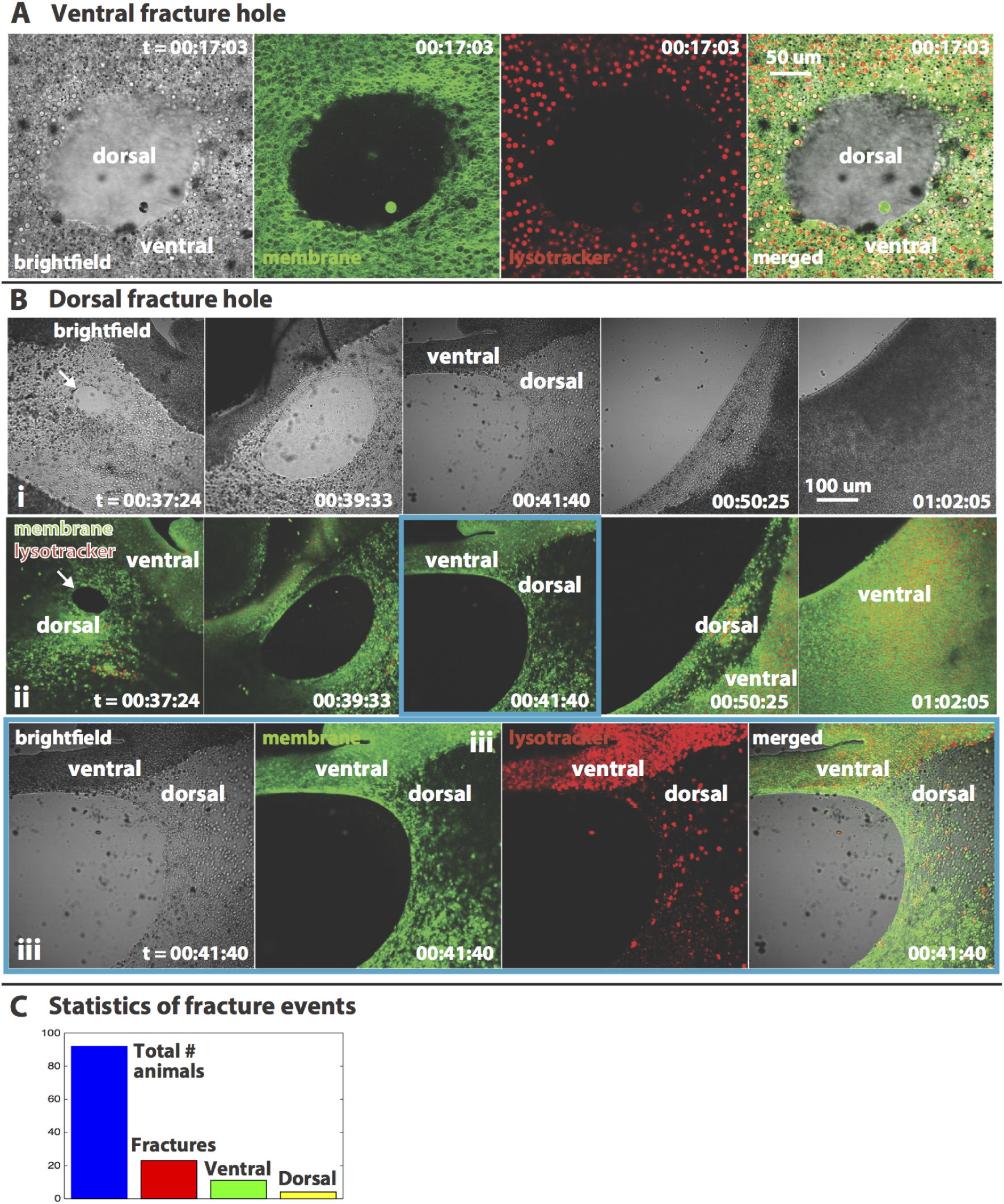
Ventral and dorsal epithelial fractures in *T. adhaerens*. (A) Ventral hole: All imaging channels corresponding to Fig. 2B (iv) are shown separately (cell membrane: green, acidic granules in lipophil cells: red). The bright-field channel reveals that the dorsal layer is still intact. (B) Dorsal hole: a) The brightfield channel corresponding to the dataset shown in Fig. 2D and the row b). c) All the imaging channels corresponding to b(iii) are shown separately (cell membrane: green, acidic granules in lipophil cells: red). The dorsal fractures are through holes inside the bulk tissue of the animals. (C) Statistics of fracture events (lower bound): Total number of animals of size ∼2-4 mm imaged in confocal microscopy experiments: N = 92, The number of fracture events: N = 42, The number of ventral fractures that healed: N = 11, The number of ventral fractures that led to dorsal fractures: N = 4.

**Extended Data Figure 2:**
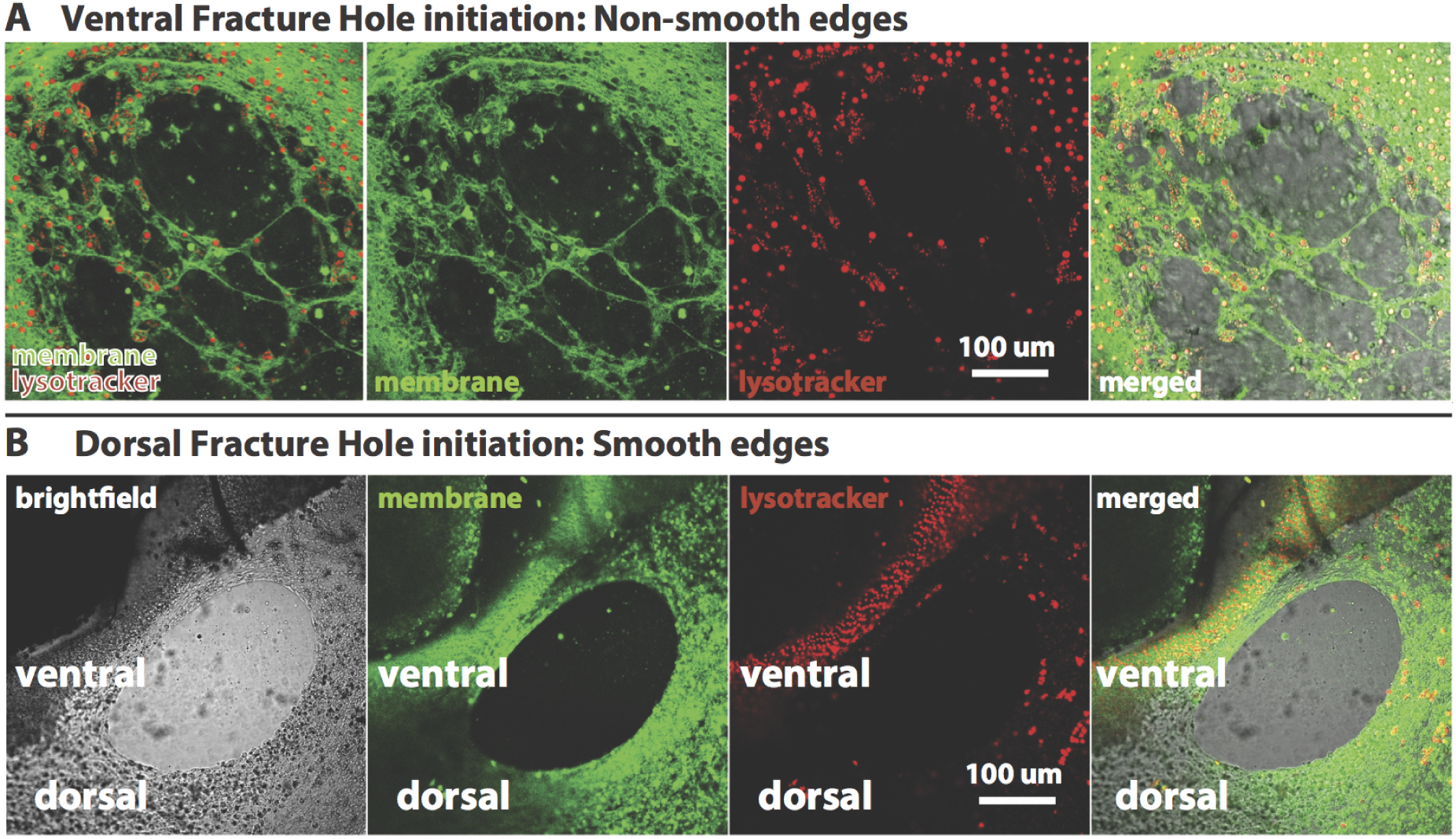
Ventral and dorsal fracture hole initiation and time dynamics. (A) Ventral fracture holes initiate as micro-fractures that coalesce to form larger holes which are brittle-like with jagged edges and broken cellular fragments (The different imaging channels are shown separately). (B) Dorsal fracture holes initiate as circular holes which grow in size and remain smooth, indicative of a more ductile dorsal tissue (The different imaging channels are shown separately to facilitate).

**Extended Data Figure 3:**
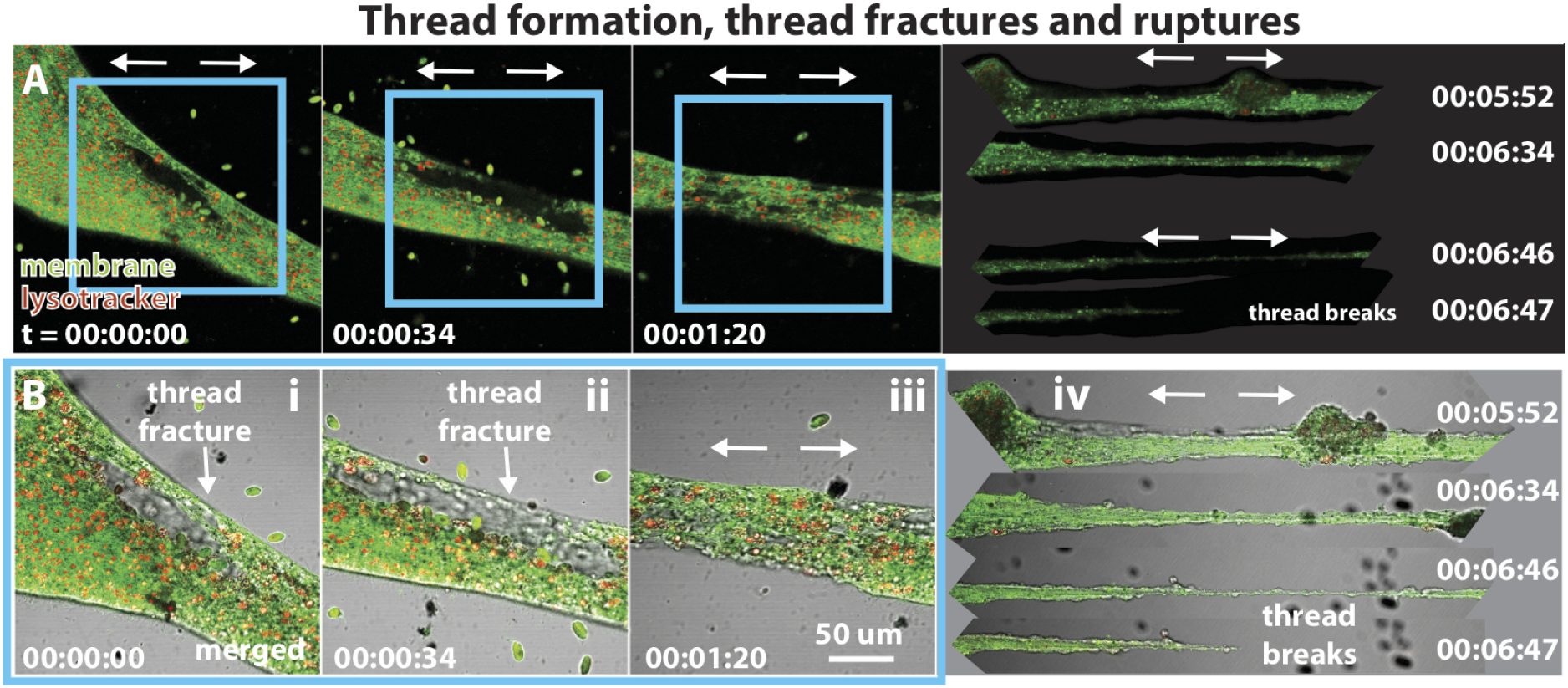
Thread formation, thread fractures and ruptures: (A) The stretched tissue between two regions of an animal that are pulling apart yield in a ductile fashion and form thread-like regions by undergoing local ‘thread fractures’. Here, the subtle difference is in the location of fractures; ventral and dorsal epithelial fractures typically occur in the bulk, whereas these thread fractures occur on the edges. Functionally, these thread fractures are cumulative over time, resulting in rapid reduction (∼few min) in the cross sectional area and lead to extremely thin (few cell layers thick) threads, which ‘rupture’ (∼few minutes) when pulled apart continuously. This mechanism plays an important role in the asexual reproduction process in these animals (Fig. 1B). (B) zoom-in versions of (A), with additional brightfield channel overlay on the images.

**Extended Data Figure 4:**
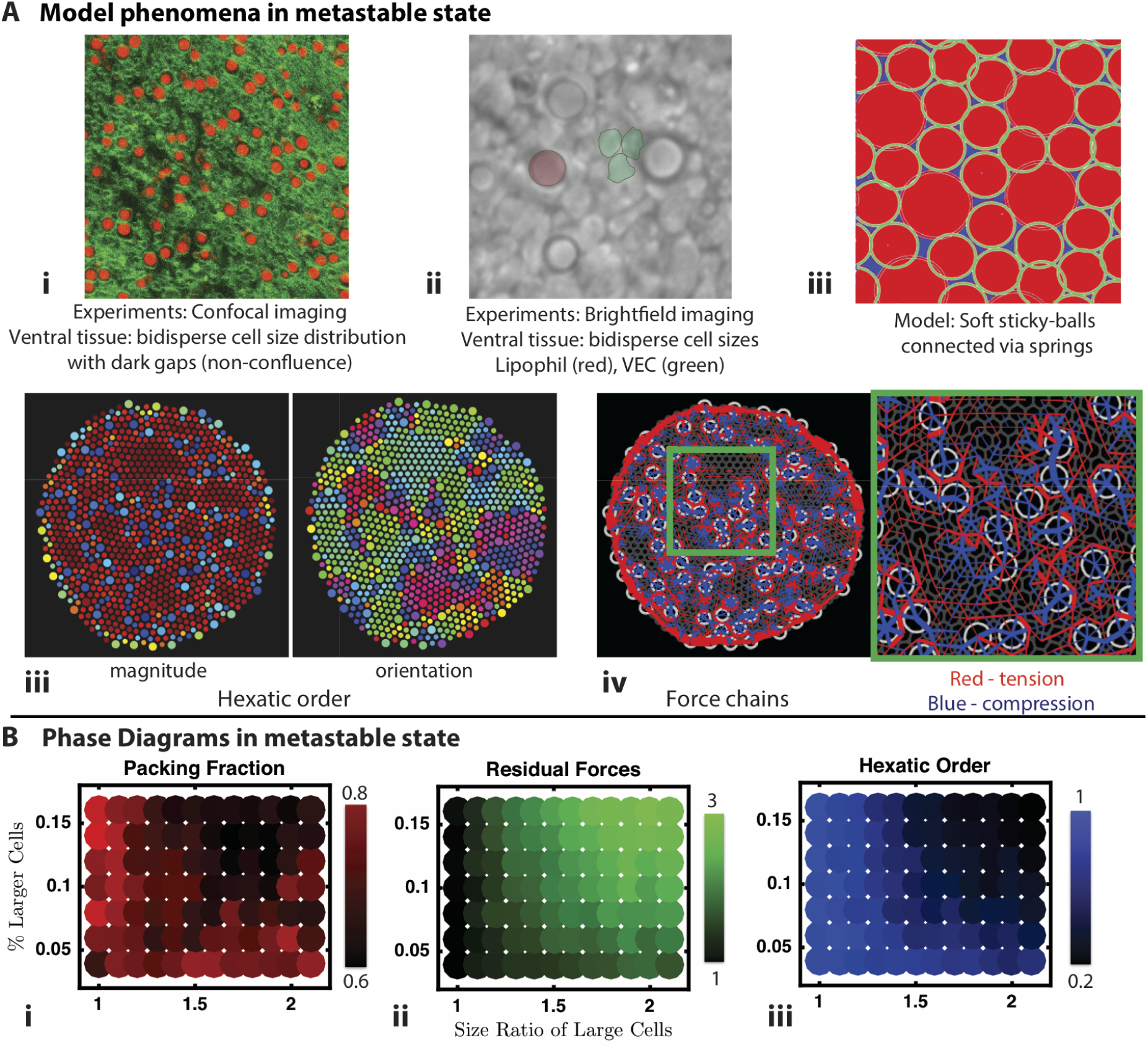
A heuristic in-silico model consistent with experimental observations. **(A) Model in metastable state:** (i) Cells within the ventral layer are a bi-disperse mixture of ventral epithelial cells and lipophil cells. The tissue is also not perfectly confluent with gaps at the ventral surface. (ii) Digital zoom of brightfield 1 *µ* m deep from the ventral surface shows bi-dispersity of cell sizes and shapes (lipophil cell - false color red, and ventral epithelial cells (VEC) - green). (iii) Close up of a quenched state of the sticky-ball model where packing fraction is calculated with a simple Monte Carlo solver using blue color regions between cells. (iii) Hexatic order - structural order measurement that reveals short range correlations of hexatic orientation for this bi-disperse system. (iv) Residual forces represented spatially through force chains demonstrates rich spatial heterogenity at the metastable state. Thickness of force chains corresponds to magnitude with red representing force under tension and blue force under compression. **(B) Phase diagrams in metastable state:** We explore a 2D cross-section of the model’s parameter space with three diagnostic quantities: (i) 1 minus the packing fraction. (ii) Residual forces, and (iii) Hexatic order. These phase spaces demonstrate how the metastable state properties can be tuned across a wide range of configurations through mixing ratios of size and percentage in an epithelial alloy.

**Extended Data Figure 5:**
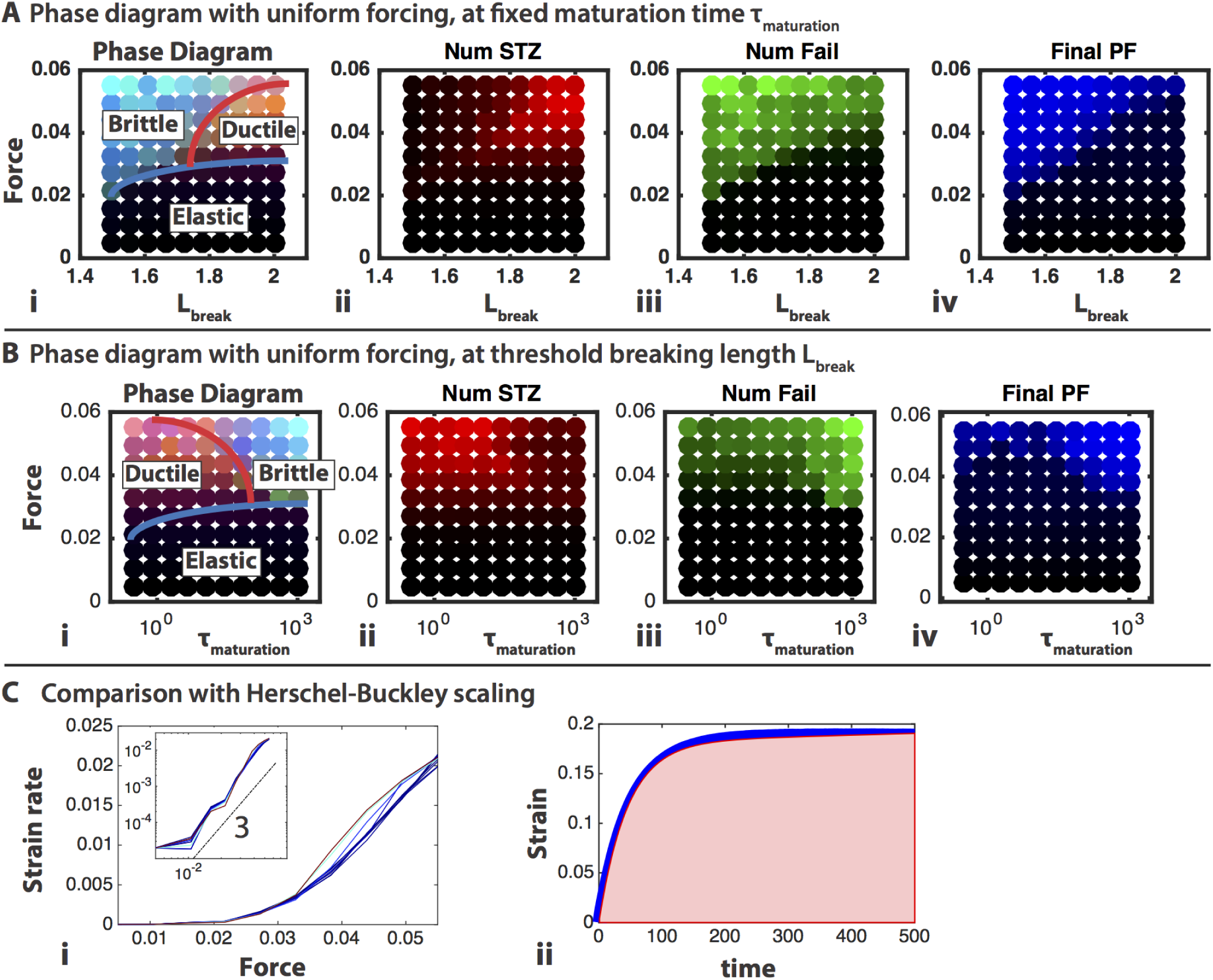
Phase diagrams with uniform forcing. Pulling the heuristic model out of equilibrium for given parameters reveals two distinct behaviors: (i) a direct yielding through new edge formation and (ii) an intermediate yielding through reshuffling of the cell-cell junction network via higher order STZs. (A) First, we consider the parameter of the maximum junctional length before failure. At small *ℓ*_*break*_ the model yields to new edge formation at smaller loading. However, at a crossover ∼*ℓ*_*break*_ = 1.7, for low loading gradients, the dominant yielding occurs through coordinated neighbor exchanges similar to STZs. This crossover value can be understood as the edge forming energy exceeding the average STZ energy barrier (which is independent of *ℓ*_*break*_). The subsequent three panels in (A)ii-iv, are each one channel of the RGB parameter space displayed on the left. From left, red is the number of STZ transformations (or CCC), green is the number of edge forming transformations (or non-CCC), and blue is a helpful diagnostic, the packing fraction, for identifying holes within the tissue (light blue is low packing). (B) Considering model tissues at different maturation times reveals another crossover between a more ductile and a more brittle behavior under distributed driving. At short maturation times, the formation of a new cell-cell junction is nearly instantaneous, so in a regime of *ℓ*_*break*_ = 1.95, the yielding is driven through STZ like (or CCC) transformations. However, when the maturation time reaches ∼10*x* the dynamical timescale, the cell-cell junctions are too slow to reform and the tissue yields through new-edge formation and holes. This rapid de-lamination is can be seen through the split channels (B)ii-iv in the reduction of STZs, the increase in new edges and the reduction of the packing fraction. (C) To study the steady-state dependence of the model’s Herschel-Buckley scaling on *τ*_*mature*_, we complete a series of fits (ii) in the extension versus time curve. The asymptotic value of this fit represents the long-time behavior of this yielding rate. Plotting the strain rate at steady-state versus the applied stress reveals a Herschel-Buckley like relationship with a finite yield stress and a 3 scaling behavior. Over the range explored *τ*_*mature*_ has only limited impact on the flow behavior of this tissue.

**Extended Data Figure 6:**
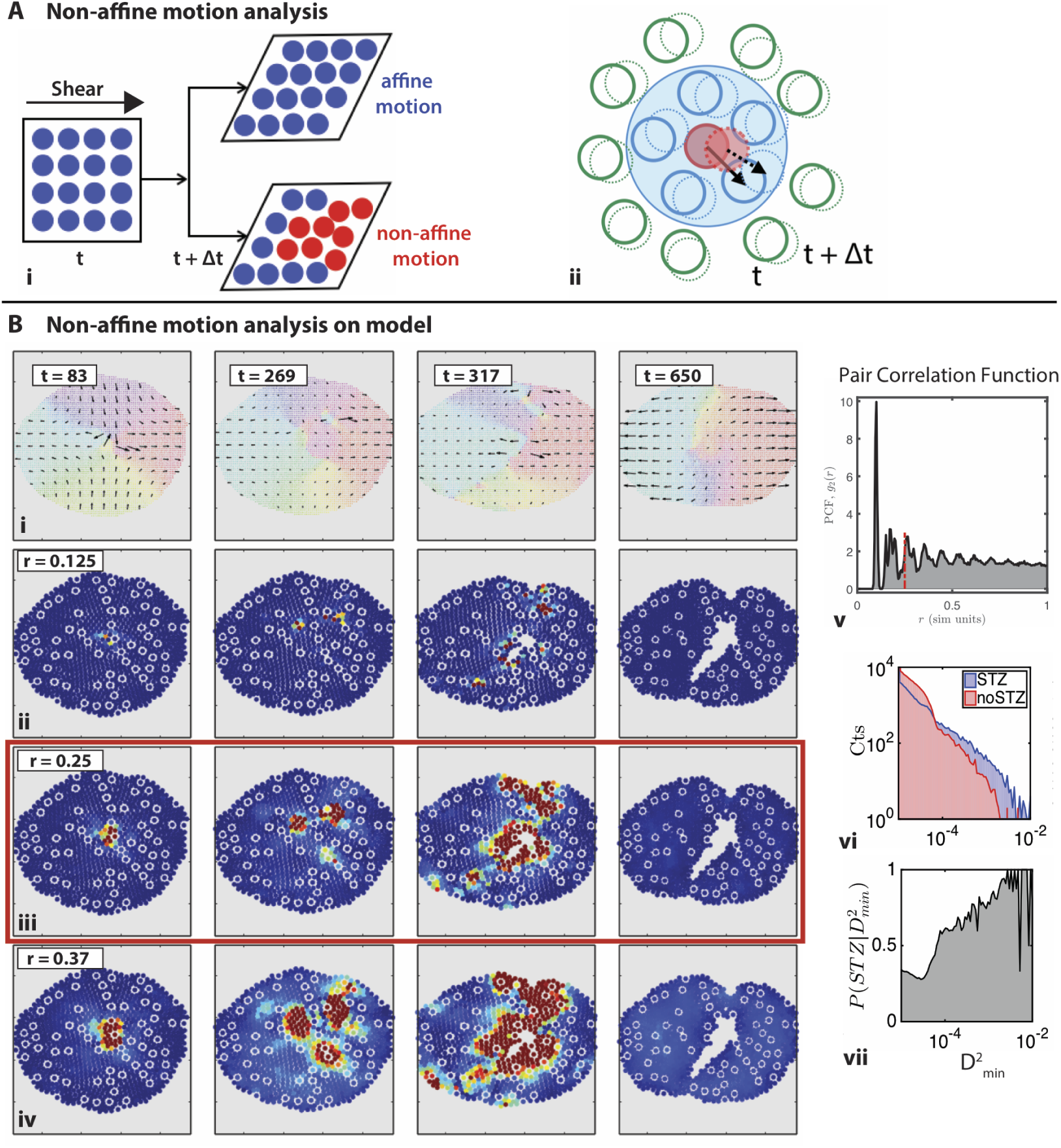
Non-affine motion analysis. Leveraging the knowledge of the microstate in numerics to develop techniques for learning about the material properties of real tissue from experimental signatures. (A) We employ a measure of non-affine motion called 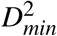 ^53^. This measure subtracts actual motion of particles from the local affine transformation inferred from its neighbors motion (ii). Large values of disagreement between actual motion and the affine projection are signaled in red (i). (B)(i) We display the strain rate field calculated from interpolation of the displacements into a grid. Color corresponds to orientation angle of the vector ranging from 0 to 2*π*. (B)(v) displays the pair correlation function showing rapid attenuation of the characteristic spacing over distances greater than 0.4 simulation units. The red dotted line signals the neighborhood size used to calculate the metric in Fig. 3. Panels (B)(ii-iv) demonstrate the effect of the neighborhood size with with 50% variation on either side of that shown in the figure (B)(iii). Above is 50% smaller neighborhood and below is 50% larger. The trade-off between sensitivity and spatial resolution is balanced by choosing the middle values B(iii). (B)(vi) shows the map between known simulation STZs and the measured 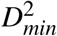. B(vii) We find that this measure is a nice diagnostic for otherwise un-observable motions with 80% of the observed large 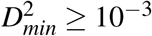 corresponding to a known neighborhood exchange via a STZ-like transformation.

**Extended Data Figure 7:**
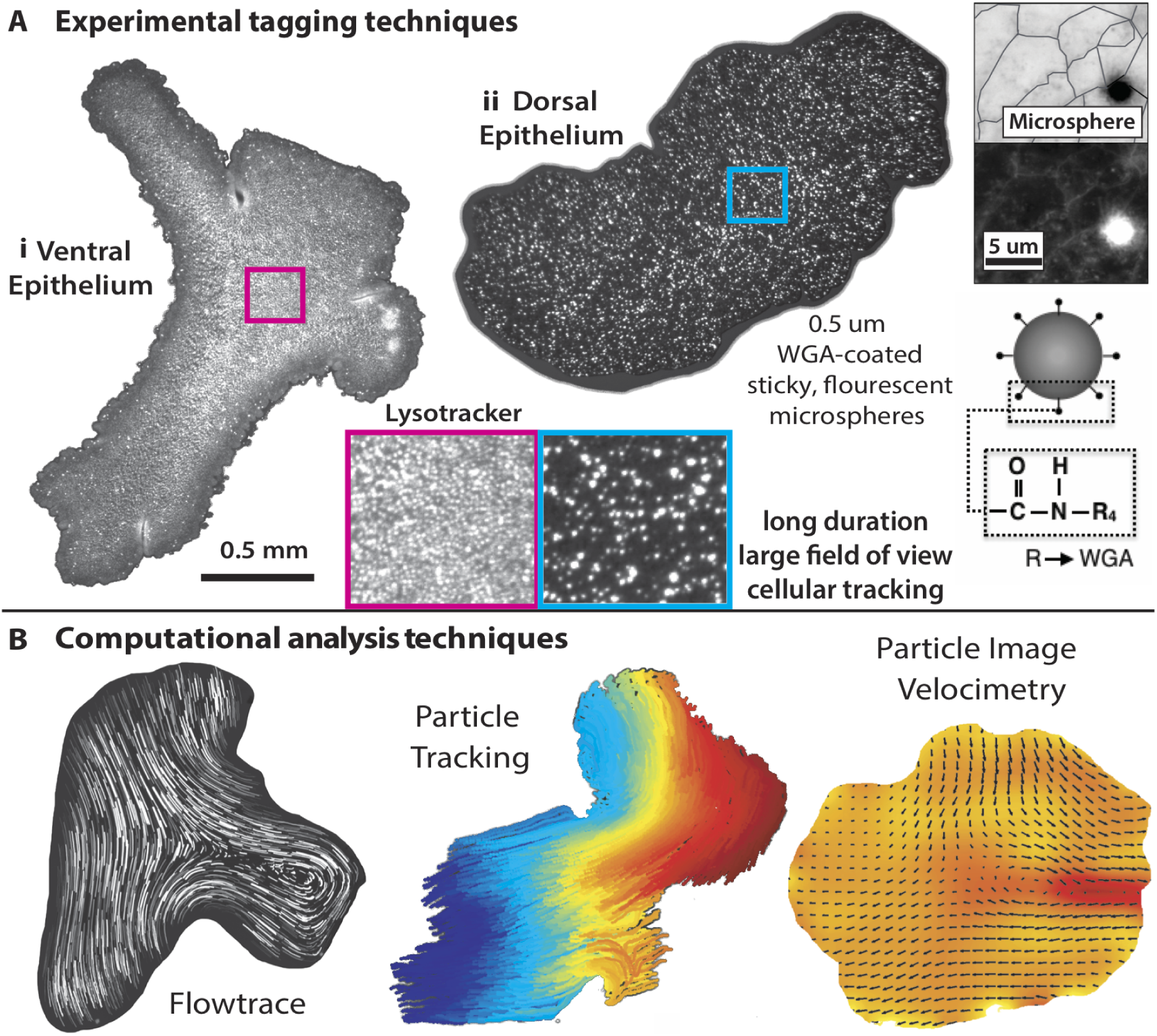
Experimental and computational techniques. (A) Experimental tagging techniques: (i) Ventral epithelium - A fluorescent lysotracker dye stains the acidic granules present in the lipophil cells, and provides a dense tagging of the entire ventral epithelium at large fields of view (∼3 mm) (Methods). (ii) Dorsal epithelium - We developed an assay to coat the surface of the epithelium with sticky fluorescent microspheres / microbeads. This provides a coarse-grained tagging (1 bead per 8 cells) of the entire dorsal epithelium at large fields of view (∼6 mm) and enables high-speed (10 fps) and long duration (∼1-5 hours) imaging (Methods). Right Insets display control experiments demonstrating that microspheres bind on cell membrane. (B) Computational data analysis techniques: We employ Flowtrace^65^ for visualization of particle trajectories, Particle tracking for quantitative non-affine motion analysis, and Particle Image Velocimetry to measure velocity fields and internal strain rate.

**Extended Data Figure 8:**
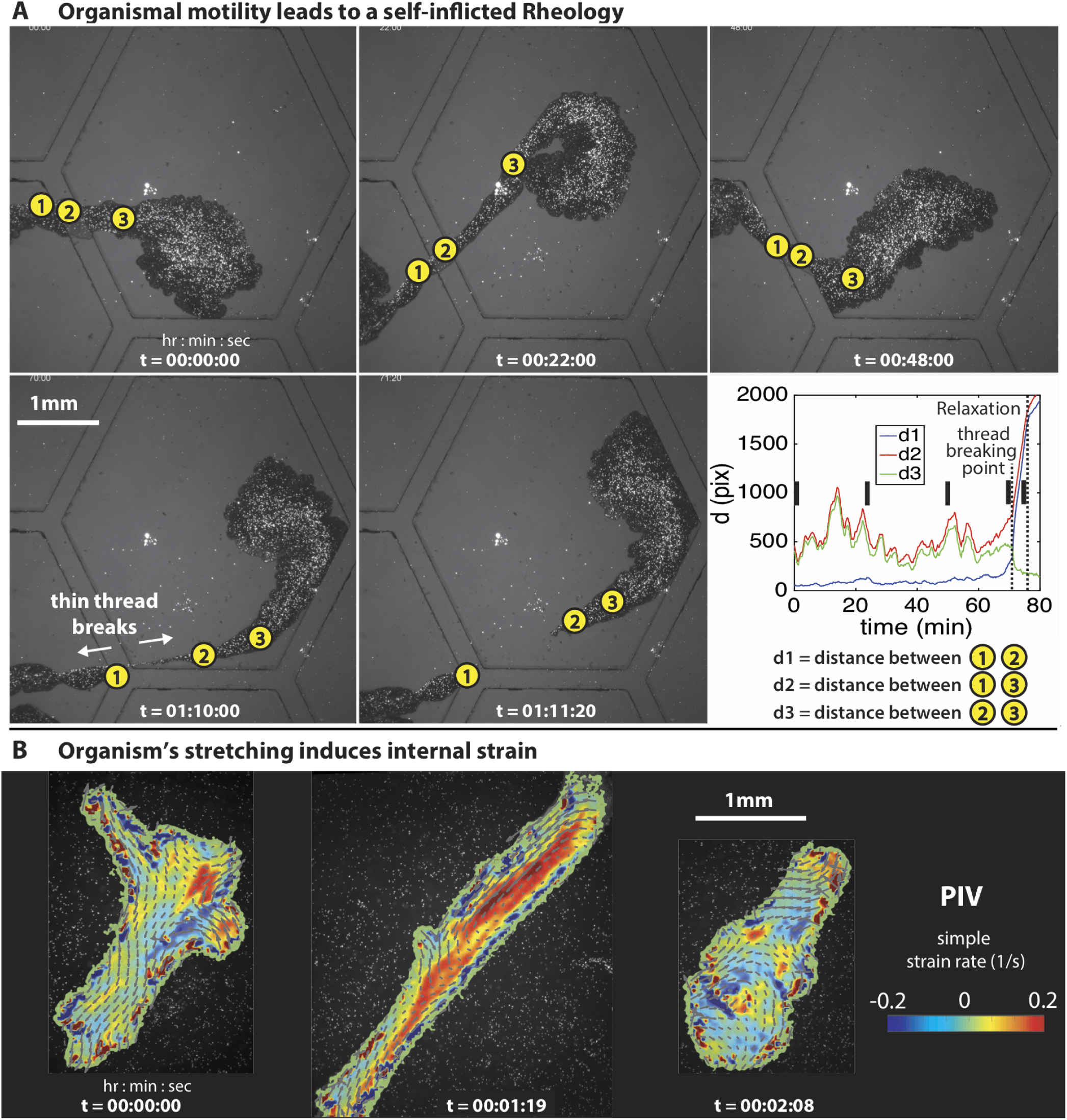
Organismal motility leads to a self-inflicted rheology in T. adhaerens. (A) Time lapse images from large field of view imaging of dorsal epithelium using the sticky microbeads assay (Methods). We plot distances between three microbeads tracked over time (lower right panel) which reveals continuous fluctuations indicative of varying loads. We also observe here the time-scales involved in the thread breaking and tissue relaxation; the black markers indicate timepoints corresponding to the displayed snapshots. (B) Time lapse images from large field of view imaging of ventral epithelium using lysotracker dye. The organism’s stretching induces peak internal strain rates (computed using a PIV analysis, Methods).

**Extended Data Figure 9:**
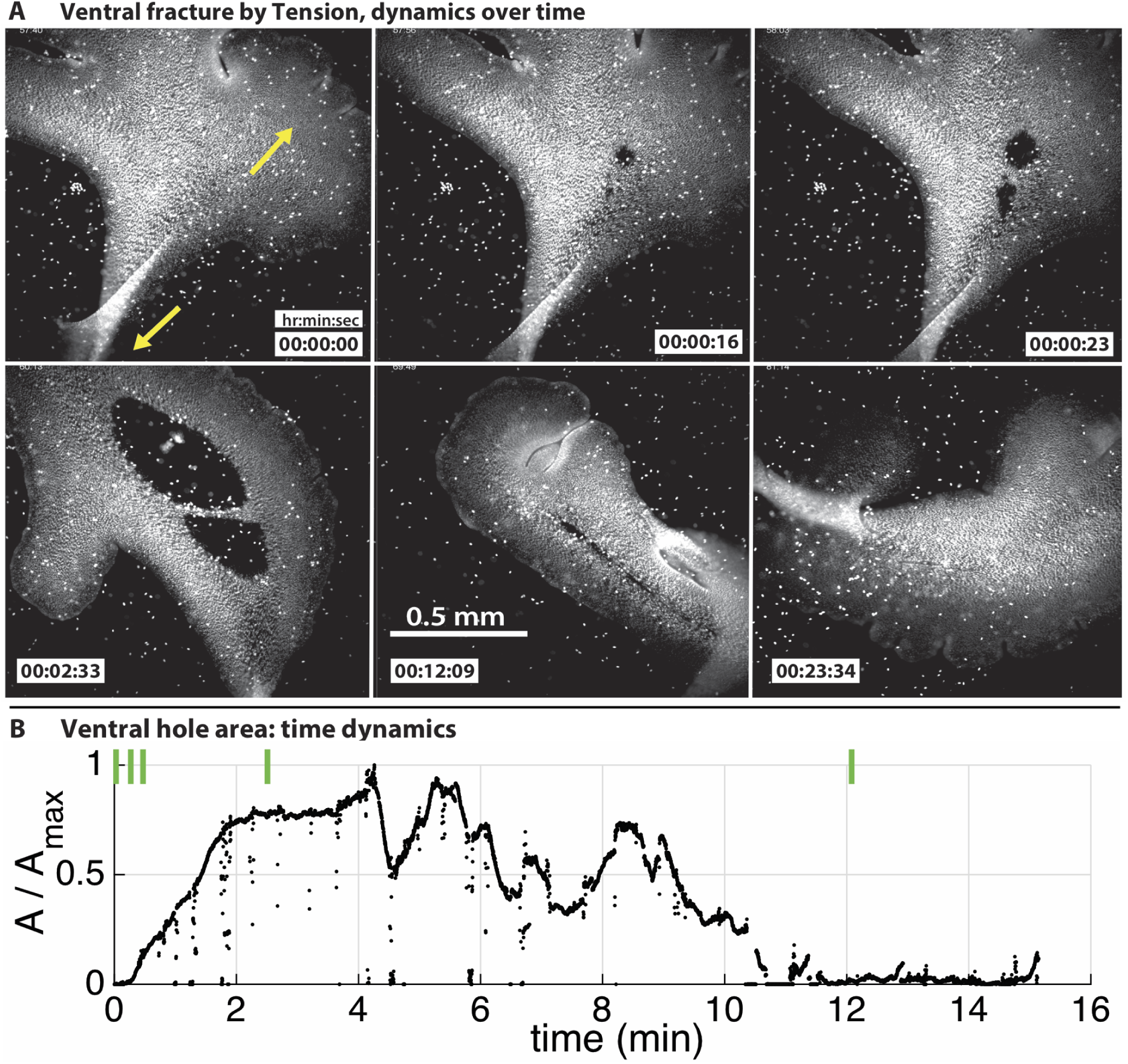
Experimental time lapse images of tensile fractures. Here, we display more information on the same dataset as in Fig. 4(A). (A) Time-lapse images showing the evolution of tensile-induced ventral fractures and their healing over a period of ∼25 mins. (B) Time-series plot of the fracture hole area in (A). The tension-induced fracture is an extremely fast process, with the hole reaching its maximum area in ∼2 mins. The healing process is completed at about ∼12 mins. The green markers indicate timepoints corresponding to the snapshots displayed above.

