## Supplementary File for "Motility induced fracture reveals a ductile to brittle crossover in the epithelial tissues of a simple animal"

#### Methods used for Images in Figures

Some experimental images of the *T. adhaerens* have been edited to remove the background, leaving only the animal for clarity. In these cases, a uniform black or white background has been introduced. Some images were edited to adjust levels of brightness, contrast and color for clarity.

#### Interpretation and formulation of the cell-cell junction bonding/debonding criterion

We relate our simple model to existing literature and ideas before jumping into the debinding criterion. The debonding/bonding criteria is central to understanding the dynamics of tissues under load and how they dynamically reconfigure (or not) under external driving.

A very interesting and useful low order expansion of the energy of a confluent tissue within a plane is captured by a elegant simple energy function applied to vertex degrees of freedom (the energy function is reminiscent of a foam)[1]:

$$\varepsilon_i = k_A (A_i - A_o)^2 + \Gamma P_i^2 + \gamma P_i$$

The second order term in perimeter (behaving like an elasticity of the cortex) is important for stabilizing the shape of the cells when the first order term (like the cell surface tension) goes negative. Without this stabilizing elasticity, cells with negative effective surface tension would stretch in an unstable fashion to infinitely thin cells with large surface area, but near $A_o$ area.[2,3]

A negative effective surface tension makes sense when we decompose the contributions of this term into two competing first order contributions[2]. $\gamma P_i \equiv \alpha P_i^{all} - \beta P_i^{shared}$, where $\alpha$ is the contribution from the cell's internal cortical tension and $\beta$ is the energy contribution from interaction with other sticky cells. In the case of a perfectly confluent tissue with no edges (periodic BC or infinite size), the energy contributions from both go negative when the shared energetic contributions exceed the cell's own surface tension $\alpha < \beta$. This energetic description is powerful in the limit where the kinetics of the bonds is fast compared to the timescales of interest[4,5]. This holds true in the long-timescale behavior of tissues flowing under tension[6].

On short timescales where the bond-lifetime is comparable to the dynamics, the transients begin to play an important role in the response of the tissue to forcing (internal or external). To study the dynamics of the bonds of the tissues, we can unfold the contributions of the two surface tension terms into:

$$\varepsilon_i = k_A(A_i - A_o)^2 + \Gamma P_i^2 + \alpha P_i + \sum_j \delta_{ji}\varepsilon_{junction}$$

With the complementary kinetics of the cell-cell junctions governed by the presence or absence of a bond, denoted by $\delta_{ij}$

$$\delta_{ij} = \begin{cases} 0, & \text{when bond is absent} \\ 1, & \text{when bond is present} \end{cases}$$

##### *Debonding criterion*

The first order description of the dynamics of the bonds under force turns to Bell's formulation of a first passage process over an energy barrier[7]. In this case, the dynamics of a single junction-junction bond has the lifetime,$\tau$, of:

$$\tau = \tau_o e^{\frac{E_O - r_O F}{k_B T}}$$

Where $\tau_o$ is the natural lifetime of the bond, $E_o$ is the bond energy. $r_o$ is the distance along the reaction coordinate between the bound and unbound state, $F$ is the applied force, $k_BT$ is the temperature in units of energy.

This can be inverted to be used as a rate in the following kinetic master equation for a two-state system:

$$\frac{\partial}{\partial t}P_{bound} = r_{binding}(1 - P_{bound}) - r_{unbinding}(F)P_{bound}$$

However, a cell-cell junction is representative of a large ensemble of bonds all working in concert to keep the cells stuck together (with higher order cis and trans assemblies forming over a hierarchy of timescales)[8,9].

We can justify approximating these ensemble dynamics as a threshold by considering distributed load amplification. When a single bond-fails, the neighbors feel a sharp increase in their load causing them to fail, and so on. We get an avalanche of bond failures that looks a lot like a threshold.

Using Bell's rate to solve for the steady state of the distribution:

$$\frac{\partial}{\partial t}P = 0 = r_b(1 - P_{ss}) - r_u e^{\beta(E_o - r_o F)}P_{ss}$$

We get a steady state probability which is dependent upon force.

$$P_{ss} = \frac{1}{1 + \frac{r_u}{r_b}e^{-\beta(E_o - r_o F)}}$$

Then the ensemble becomes a Bernoulli distribution if the binding events are independent:

$$p(n; N(t), P(F)) = (P_{ss}(F))^n (1 - P_{ss}(F))^{N(t)-n}$$

Where $n$ is the number of bound junctions. $N(t)$ is the recruitment of cis-bonds which create these large scale islands of cell-cell adhesion clusters with a timescale in the $\sim 10$ of seconds[9]. $P_{ss}(F)$ is the steady state which is determined by a single external force distributed over the population of attached bonds.

Recall that the mean of a Bernoulli distribution is simply $\langle n \rangle = N(t)P_{ss}$

Next, we calculate for a given $F_{pull}$ the mean force distributed to each of the bonds[10]. This takes for the form of:

$$\langle F \rangle = \frac{F_{pull}}{\langle n \rangle}$$

We can take the analytical form of the expectation value for a Bernoulli random process:

$$\langle F \rangle = \frac{F_{pull}}{N(t)P_{ss}(\langle F \rangle)} \qquad = \frac{F_{pull}\left(1 + \frac{r_u}{r_b} e^{-\beta(E_o - r_o\langle F \rangle)}\right)}{N(t)}$$

The calculation does not have an analytical solution:

$$N(t)\langle F \rangle - \frac{r_u}{r_b} e^{-\beta(E_o - r_o\langle F \rangle)} = F_{pull}$$

So given that this function set transcends algebraic interrogation, we expanded our exponential as a Taylor series to $\mathscr{O}(F)$. This gives us the result:

$$\langle F \rangle \sim \frac{\left(1 + \frac{r_u}{r_b} e^{-\beta E_o}\right) F_{pull}}{N(t) - \frac{r_u}{r_b}\beta r_o e^{-\beta E_o} F_{pull}}$$

This is the low order approximation of the force dependence and looks like the functional form:

$$\langle F \rangle(F_{pull}) \sim \frac{C_1}{\frac{1}{F_{pull}} - C_2}$$

The average force on each two-state system diverges when $N(t)F_{pull}^{-1} = \frac{r_u}{r_b}\beta r_o e^{-\beta E_o}$.

If we recall that our probability of being bound looks like:

$$P_{ss} = \frac{1}{1 + \frac{r_u}{r_b} e^{-\beta(E_o - r_o\langle F \rangle)}}$$

Then, we can plug in our calculated mean force with feedback to give us the probability of steady state binding for a given pull force:

$$P_{ss}(F_{pull}) \approx \frac{1}{1+\frac{r_u}{r_b} e^{-\beta\left(E_o - \frac{r_o C_1}{F_{pull}^{-1} - C_2}\right)}}$$

As $F_{pull} \to C_2^{-1}$ the exponential diverges rapidly sending the binding probability toward 0. For sufficiently high affinities, this function is well approximated by a Heaviside step function.

$$P_{ss}(F_{pull}) \approx P_{ss}(\langle F \rangle = 0)\left(1 - \Theta_{Heaviside}(F_{pull} - F_c)\right)$$

The distribution, and therefore the probability of debonding, is then captured by a single value, the force threshold of debonding, $F_c$. In the elastic limit, we can translate our threshold yield force, into a geometric criterion allowing us to port over efficient methods in computational geometry to quickly update the connectivity at each time-point.

##### *Comparison of debonding criteria with results from contact mechanics*

This single threshold approach is complemented by the formulation of Hertz contact mechanics with adhesion energy formulated by Johnson and coworkers[11]. The JKR theory for contact mechanics asks the question about the interplay between the stored elastic energy of deformation and the surface energy of contact. The math is identical to fracture mechanics in that the critical values are tight thresholds determined by the radius, the interaction energy, and the critical load to separate the two surfaces. In this picture, strain and stress are related through the modulus and there is no transients. From the perspective of JKR theory, there is a finite pull-off load or force which is equal to the crossover in the energy contributions (being attached or not). This crossover occurs at:

$$F_{\text{pull off}} = 3\pi\Delta\gamma R$$

Where $\Delta\gamma = \gamma_1 + \gamma_2 - \gamma_{12}$.

Notice that the energy between the interfaces is the integral of the LJ interaction potential to give the work to remove the sphere. While the interpretation is slightly different, the debonding threshold comes out similarly from this very distinct approach. [cite same as above]

##### *Bonding criterion*

For rebonding We propose a single timescale for maturation, $\tau_{mature}$, on the 'stiffness' of the cell-cell junction. This single timescale is consistent with the local recruitment of the cadherin complex, which we have represented above as $N(t)$. The number of cytoskeletal links between the two cells correlates to the stiffness and the number of links, and is determined by the number of cadhearins in the local cluster.

With $N(t) = N_o(1 - e^{-t/\tau_{mature}})$, the cell-cell bond energy looks like:

$$\varepsilon_{bond}(t) = N_o \varepsilon_{junction} \left(1 - e^{-t/\tau_{mature}}\right)$$

Where $t$ is the age of the bond since entering into the triangulation and the geometric cutoff, $L_{break}$.

By focusing our attention on the behavior of the connectivity matrix to emphasize the dynamics of cell-cell junctions over cellular shape dynamics, we open up the door to a careful comparative study incorporating higher order cell dynamics more consistent with recent work[12,13]. Undoubtedly cell shape will contribute to our understanding of the collective action in a tissues response to fast timescale forcing and will be a target in the next generation of models.

##### *Applying bonding/debonding criteria to low order dynamics of a model tissue*

Following above, we wrote two models to explore the impact of cell shape [results not presented in detail here]. One where cells are promoted to a large number of degrees of freedom and their shape is governed by

$$\varepsilon_i = k_A(A_i - A_o)^2 + \Gamma P_i^2 + \alpha P_i + \sum_j \delta_{ji} \varepsilon_{junction}$$

The second, we removed the shape dynamics and replaced it with a characteristic cell compliance (of the form of a Hertizian interaction). We capture qualitatively the compliance of the cells (in a two body way) by linearizing the elasticity of the vertex model energy and using those fictious springs between the cells. This simplified energy emphasizes the central role of dynamics of this connectivity network:

$$\varepsilon_i = \sum_j \delta_{ji}\varepsilon_{junction}$$

We can expand the energy of the junction term into: $\varepsilon_{junction} = \int_0^{L_{break}} d\ell \cdot \left(-k_{junction}\Delta\ell\right)$. This becomes the energy of the bond.

The dynamics of this network emerge as two simple possibilities. (1) the cell-cell junction is stretched to the threshold of failure resulting in the breaking of the bond and a formation of a new edge ('edge' meaning that we do not locally conserve the number of constraints, no CCC). (2) The network can undergo a tranformation similar to a shear transformation zone or t1 (which conserves the number of local constraints). The STZ type transformations occur when the energy of one configuration is more favorable than the other, however, the time dynamics of rebonding still control the kinetics of the bond swap. The old bond will break and the new bond will slowly form.

This competition between stress relaxation mechanisms forms the basis of the competition between fracture and flow.

### Steady state solutions to dissipative failure of a toy model

The simplest viscoelastic system comes in the form of a Maxwell element: a damping in parallel to a series system of a damping and a spring.

We can define the stress of our linear spring as $\sigma_s(t) = k\ell_s(t)$, where $\ell_s$ is the strain.

For the simple damper, the stress goes as : $\sigma_d(t) = \eta \frac{\partial}{\partial t}\ell_d(t)$.

If we take our results from up above, we suggest that a force threshold looks like a good low order model for the rupture criterion, thus we can apply the well known stress criterion for failure. This suggests that failure occurs at a value $\sigma^*$.

One can then apply a imposed strain rate to the connecting walls $\dot{\ell}_{pull}(t)$ and generate the equation for the growing internal stress as:

$$\frac{\partial}{\partial t}\sigma_s = \eta\dot{\ell}_{pull} - k\ell_s$$

This relationship has the straightforward implication that the steady state is that the material will fail if the imposed pull rate is faster than $\dot{\ell}_{pull} \geq \frac{\sigma^*}{\eta}$.

If the flowing material instead falls into the Herschel-Buckley equation of yield stress fluids, as we might expect for a flowing material, this relationship instead becomes:

$$\frac{\partial}{\partial t}\sigma_s = \eta_{HB}\dot{\ell}^n_{pull} - \sigma_y - k\ell_s$$

With $n$ becoming the power of the HB equation ($\sim 3$ for our sticky material) and $\sigma_y$ being the yield stress.

Outside the transients this means that failure will occur for imposed pulling rates: $\dot{\ell}_{pull} \geq \left(\frac{\sigma^* - \sigma_y}{\eta_{HB}}\right)^{1/n}$.

If we combine these two results, we find that the two complementary mechanisms of relaxation can combine forces to give:

$$\frac{\partial}{\partial t}\sigma_s = \eta_{HB}\dot{\ell}^n_{pull} - \sigma_y + \eta\dot{\ell}_{pull} - k\ell_s$$

At small pull rates, the relaxation timescale of the tissue dominates. At large pull rates the relaxation through neighborhood exchange takes over.

The cross-over value occurs when: $\eta_{HB}\dot{\ell}^n_{cx} - \sigma_y = \eta\dot{\ell}_{cx}$ which becomes:

$$\dot{\ell}_{cx} = \frac{\left(\frac{2}{3}\right)^{1/3\eta}}{\chi} + \frac{\chi}{2^{1/3}3^{2/3}\eta_{NB}}$$

where $\chi \equiv \left(9\eta^2_{HB}\sigma_y + \sqrt{3}\sqrt{27\eta^4_{HB}\sigma_y - 4\eta^3_{HB}\eta^3}\right)^{1/3}$

A followup to this question is: does the cross-over between subcellular and cellular scale flows occur before the threshold for fracture?

Generically, there are four outcomes to a pull experiment in this toy model. (1) At low pull rates, and fast cellular relaxation timescales, the tissue yields by sub-cellular flow alone. (2) At medium pull rates with a slow cellular relaxation relative to the rate of loading, the tissue can cross over into a flow by neighborhood rearrangements. (3) When the yield stress is higher than the fracture stress, the tissue never flows via rearrangement and only flows via subcellular processes up to breaking (on these fast timescales, the bond-lifetime controls the kinetics rearrangments) (4) At sufficiently fast loading rates, the tissue fails catastrophically as the loading cannot be compensated for via either of the available stress relaxation mechanisms (on these timescales, a third candidate might be stress induced oriented cell division)[14].

This toy model illustrates that even with higher order mechanisms of cell relaxation on short timescales[15], the concept of pitting flow versus fracture as competing mechanisms holds a useful tool for understanding. The presence of such a mechanism enriches the character of tissue under fast loading timescales and suggests an interesting interplay between flow mechanisms which can serve complementary roles.

#### Kinetic perspective on the competition between fracture and flow

The central theoretical concept in this work is the competition between edge formation and neighborhood exchange (or fracture versus flow). While in the main text, we developed an argument on the foundation of a numerical study with a simple real-space interpretation, here we attempt to formulate the argument in a more model-agnostic perspective. This toy argument is a simple tool for understanding outcomes and coming up with sharp definitions.

One possible approach to this is from the perspective of the kinetics on a dynamic energy landscape. Let's consider a three state system with the following properties.

- State 1 corresponds to a configuration of the neighborhood matrix between 4 particles. It is metastable at zero force.
- State 2 corresponds to the lowest energy transformation of that neighborhood matrix which preserves the number of local constraints. It is metastable at zero force and is an STZ like transformation away from state 1.
- State 3 corresponds to a fractured state where the number of constraints is fewer than state 1 or 2.

First, it is critical to link these multistate kinetics to the practical definition of ductile and brittle used in this text. A generically ductile material will pass from State 1 to State 2 a majority of the time. Whereas a generically brittle material will preferentially pass from State 1 to State 3. A final possibility is that material could pass quickly through State 2 en route to State 3. In this language, this material will still behave in a 'brittle' fashion, if the lifetime in State 2 is insignificant compared to the timescales of the problem. If a material has a long-lived occupancy in State 2 at any force, we call it ductile. A more ductile material will almost exclusively use the State 1 to State 2 path on this kinetic path for a large range of forcing.

Let's assume that the energy needed to generate new edge is $\varepsilon^B_{break}$. The activation energy for this transformation is the same as the bounding energy.

Let's further assume that the energy needed to undergo an STZ transition is $\varepsilon^B_{STZ}$. This is a barrier height and represents a combination of both elastic and adhesive energies at play. Noteworthy, this landscape is time dependent controlled by timescales set by $\tau_{mature}$ and the neighborhood topology which we approximated by a computational geometry problem (reasonable in the limit of stiff cells and lower adhesion energies i.e. low shape parameter).

We apply an external force to state 1 driving it toward state 2.

On the long time limit (where the transients of the bond kinetics are short compared to the timescales), The outcome of the pull (whether it STZs or breaks) will be dependent upon the relationship between the activation energy for the STZ, $\varepsilon_{STZ}$ (which is a configuration dependent calculation) versus the energy to severe the connection between cells and create new edge, $\varepsilon_{break}$.

When $\varepsilon_{break} < \varepsilon_{STZ}$, new edges will form. In the other case, the tissue will flow through STZs.

This relationship is born out in the crossover observed at low $\tau_{mature}$ between fracture and flow in the numerics. Recall that $\varepsilon_{break} \sim \ell^2_{break}$ whereas $\varepsilon_{STZ}$ will be essentially independent of the edge-formation energy and will be determined by the complicated interplay of cell compliance and local configuration. Thus we expect a cross-over in the qualitative behavior of the tissue when these energy scales intersect (in the long-time limit).

On the timescales where the cell-cell junction kinetics become relevant, the problem requires a little more careful a treatment.

By a mean first passage process, the kinetics on this energy landscape can be approximated via transition state theory using the energy barrier height separating the valleys. This is essentially a study of the rate of extreme values in energy given a finite temperature.

The rates between these three states then look something like:

$$r_{1\to2} \sim e^{-\beta\varepsilon_{STZ}} \qquad r_{1\to3} \sim e^{-\beta\varepsilon_{break}} \qquad r_{2\to3} \sim e^{-\beta\varepsilon_{23Barrier}(t_{age})}$$

There is now a new term in here which has an interesting time-dependence, $\varepsilon_{23Barrier}(t)$ which represents the barrier between the new STZ state and the state with new edges. Due to the maturation time of the cell-cell junction, this is time dependent. The junction stabilizes with time.

The energy of this relationship looks something like:

$$\varepsilon_{23Barrier}(t_{age}) \sim \varepsilon_{break} - \varepsilon_j N_o \left(1 - e^{-t_{age}/\tau_{mature}}\right)$$

Where $t_{age]}$ is acting as the age of the new cell-cell junction, just following its recent STZ type transformation.

Using a Bell-like dependence on force in the direction of a transformation, the rates will take the form:

$$r_{1\to2} \sim e^{-\beta\varepsilon_{STZ}+F\cdot\hat{r}_{12}\Delta r_{12}} \qquad r_{1\to3} \sim e^{-\beta\varepsilon_{break}+F\cdot\hat{r}_{13}\Delta r_{13}} \qquad r_{2\to3} \sim e^{-\beta\varepsilon_{23Barrier}(t_{age})+F\cdot\hat{r}_{23}\Delta r_{23}}$$

Where $F\cdot\hat{r}_{ij}\Delta r_{ij}$ is the applied force projected along the reaction coordinate between state $i$ and state $j$ multiplied by the projected distance along the reaction coordinate of the minimum.

We can use this simple back of the envelop calculation to approximate the expected probability of ending up in State 3 within a time, $t_{observe}$ and compare that to the probability of ending up in state 2 within time $t_{observe}$. This ratio gives us a quantitative measure of where on the ductile-brittle spectrum we might expect to find a model system for given $\varepsilon_{STZ}, \varepsilon_{break}, \varepsilon_j N_o, \tau_{mature}$, and $F\cdot\hat{r}_{ij}\Delta r_{ij}$.

For the case of $\tau_{mature} \to 0$, the time dynamics of the second term does away and we can just define this tractable matrix collecting all of our rates:

$$K \equiv \begin{bmatrix} -e^{-\beta \varepsilon_{STZ}+F} - e^{-\beta \varepsilon_{break}+F} & e^{-\beta \varepsilon_{STZ}+F} & e^{-\beta \varepsilon_{break}+F} \\ 0 & -e^{-\beta \varepsilon_{break}+F} & e^{-\beta \varepsilon_{break}+F} \\ 0 & 0 & 1 \end{bmatrix}$$

The measure of the ratio of trajectories which pass through state 2 to state 3 tells us something about where this failure lies on the ductile-to-brittle spectrum. In the simplest one step, this looks like:

$$\frac{\text{number of STZ}}{\text{number of fracture}} = \frac{e^{-\beta \varepsilon_{STZ}}}{e^{-\beta \varepsilon_{break}}}$$

.

In the first step the ductility of the flow-fracture material looks like something which is dependent upon only the energy barriers.

But this doesn't catch the intuition that the lifetime in the second state is also important for understanding the material response at the threshold of failure. We can learn something about that by evolving the probabilities forward in time for a small period $t_{observe}$.

After time $t_{observe}$ the dynamics will have gone:

$$|P(t = t_{observe}\rangle = K^{t_{observe}} |P(t = 0)\rangle$$

To generalize taking this matrix to the $t_{observe}$ power, we can find the transformation matrix which diagonalizes it and then take the power of the diagonalized matrix sandwiched by the tranformation:

$$K^{t_{observe}} = S \Lambda^{t_{observe}} S^{-1}$$

This equivalent to finding the normal modes of this kinetic matrix, and using the time evolution operator to advance forward.

Which we can then apply to our initial state:

$$|P(t=0)\rangle = \begin{bmatrix} 1 \\ 0 \\ 0 \end{bmatrix}$$

We calculate our a template matrix raised to the power of $n \to t_{observe}$ such that $a \to e^{-\beta \varepsilon_{STZ}+F}$ and $b \to a \to e^{-\beta \varepsilon_{break}+F}$

$$\begin{bmatrix} -a-b & a & b \\ 0 & -b & b \\ 0 & 0 & 1 \end{bmatrix}^n = \begin{bmatrix} (-a-b)^n & (-b)^n - (-a-b)^n & \frac{(-b)^{n+1}+b}{b+1} \\ 0 & (-b)^n & \frac{(-b)^{n+1}+b}{b+1} \\ 0 & 0 & 1 \end{bmatrix}$$

The outcome of this calculation will take the ratio of the probabilities of being in state 2 and that of state 3. For small values of this ratio, the system appears brittle and for large values of this ratio, the system appears more ductile. The pathway from state 1 to 3 will define it dissipation.

The next step is to consider how a system with a finite maturation time behaves under force. This is trickier, but can be approximated in a couple ways. One of which is by adding a long chain of states which keep track of the age's effect on the transition probability. For the purpose of this work, it is sufficient to play out the thought experiment: if the mean rate from 2 to 3 is greater then the material will behave in a more brittle manner with a finite observation time. This means that an order 1 maturation time will have the effect of driving the system to more brittle-like behavior.

Clearly, this toy representation is an oversimplification of the rich spatio temporal dynamics of yielding. It neglects the coupling and facilitated dynamics whereby an STZ can initiate another STZ nearby coupled via the 2D shear transformation Green's function[16]. It is also worth cautioning that this toy model is a configuration dependent oversimplification. There are many configuration dependent energies that can be calculated for different types of this problem (e.g. any of the energy functions suggested in the earlier paper are great candidates), but the point is that they are strongly local configuration dependent and thus that is what makes cell-resolved modeling important for understanding failure in these heterogenous materials at wavelengths comparable to the constituent cell size.

A final note is that the statistics show here assume that thermal-like (Boltzmann distributed) flucuations are driving these transitions. Since these flucuations are active in nature, other distributions may be better descriptions to understand the finer points.

Despite its extreme simplicity, this second toy representation helps us communicate the central role between flow and fracture by making clear definitions, reasonable calculations and a playground for exploring the implication of input parameters on the position in the ductile to brittle spectrum of response.

### Supplementary Information: Videos

1. **Video1.mov: Asexual reproduction by fission in *Trichoplax adhaerens*.**
   Time-lapse quasi-dark field imaging of animals in lab culture conditions using a DSLR camera. A single animal ‘splits into two’ by a binary fission process in about one hour. Video playback is sped up, and time stamp represents hours and minutes. Scale bar: 3 mm.

2. **Video2.mov: Physiological tissue fractures in the ventral epithelium of *Trichoplax adhaerens*.**
   Time-lapse quasi-dark field imaging of animals in lab culture conditions using a DSLR camera. The ventral epithelium sustains fracture holes which heal completely in about one hour. Video playback is sped up, and time stamp represents hours and minutes. Scale bar: 1 mm.

3. **Video3.mov: Physiological tissue fractures in the dorsal epithelium of *Trichoplax adhaerens*.**
   Time-lapse quasi-dark field imaging of animals in lab culture conditions using a DSLR camera. The dorsal epithelium sustains fracture holes which grow in size and do not heal. These animals eventually become long string-like animals over about 7 hours. Video playback is sped up, and time stamp represents hours and minutes. Scale bar: 3 mm.

4. **Video4.mov: Ventral tissue fractures at a cellular resolution.**
   Time-lapse confocal imaging of animals in an open dish configuration matching native culture conditions. The ventral epithelium is tagged with a fluorescent cell membrane dye (green), and a lysotracker dye (red) that labels acidic granules in lipophil cells. Videos are looped over 5 secs, and the time stamps on images represent minutes and seconds. Scale bar: 50 um.

5. **Video5.mov: Model results with steady pulling.**
   We display the phase diagram from our heuristic tissue model, which explores a parameter sweep of steady force gradient versus the threshold strain for breaking cell-cell bonds. Next, we sequentially show simulations that demonstrate cases of elastic, ductile and brittle tissue properties — and their corresponding parameters on the phase diagram. The time stamps represent simulation units.

6. **Video6.mov: Model results with unsteady pulling.**

We show simulations with unsteady pulling of the model tissue, and demonstrate how this captures both fractures and healing. The time stamps represent simulation units.

7. **Video7.mov: Tension force-induced brittle fracture in *Trichoplax adhaerens*.**
   Time-lapse fluorescence microscopy imaging reveals a tensile force-induced fracture. The ventral epithelium is tagged using a lysotracker dye, which labels acidic granules in lipophil cells. Video playback is real time, and time stamp represents minutes and seconds. Scale bar: 0.5 mm.

8. **Video8.mov: Shear force-induced brittle fracture in *Trichoplax adhaerens*.**
   Time-lapse fluorescence and bright field microscopy imaging reveals a shear force-induced fracture in the ventral epithelium. The dorsal epithelium is tagged using 0.5 um sticky, fluorescent micro-beads. A Particle Image Velocimetry (PIV) analysis is carried out to quantify the internal tissue velocity fields (highlighted by green arrows). Video playback is sped up, and time stamp represents minutes and seconds. Scale bar: 1 mm.

9. **Video9.mov: Non-affine motion analysis on experimental data.**
   Time-lapse fluorescence and bright field microscopy imaging reveals a shear force-induced fracture in the ventral epithelium. The dorsal epithelium is tagged using 0.5 um sticky, fluorescent micro-beads. A Particle Tracking analysis is carried out to quantify the non-affine motion of the micro-beads (with magnitude highlighted by colors). Video playback is sped up, and time stamp represents minutes and seconds. Scale bar: 1 mm.

10. **Video10.mov: Correlation between non-affine motion and internal strain rate, in experimental data.**
   Time-lapse fluorescence and bright field microscopy imaging reveals a shear force-induced fracture in the ventral epithelium. The dorsal epithelium is tagged using 0.5 um sticky, fluorescent micro-beads. Larger values of non-affine motion are thresholded and overlayed (white dots) on contours (jet colorbar) of the internal strain rate calculated from the PIV analysis. Video playback is sped up, and time stamp represents minutes and seconds. Scale bar: 1 mm.
